# Genomic basis and evolutionary potential for extreme drought adaptation in *Arabidopsis thaliana*

**DOI:** 10.1101/118067

**Authors:** Moises Exposito-Alonso, François Vasseur, Wei Ding, George Wang, Hernán A. Burbano, Detlef Weigel

**Author notes:** Current address: CNRS, UMR5175, Centre d’Ecologie Fonctionelle et Evolutive, F-34000 Montpellier, France (FV). Current address: Computomics, Davis, California, United States (GW).

## Abstract

Because earth is currently experiencing unprecedented climate change, it is important to predict how species will respond to it. However, geographically-explicit predictive studies frequently ignore that species are comprised of genetically diverse individuals that can vary in their degree of adaptation to extreme local environments; properties that will determine the species’ ability to withstand climate change. Because an increase in extreme drought events is expected to challenge plant communities with global warming, we carried out a greenhouse experiment to investigate which genetic variants predict surviving an extreme drought event and how those variants are distributed across Eurasian *Arabidopsis thaliana* individuals. Genetic variants conferring higher drought survival showed signatures of polygenic adaptation, and were more frequently found in Mediterranean and Scandinavian regions. Using geoenvironmental models, we predicted that Central European populations might lag behind in adaptation by the end of the 21^st^ century. Further analyses showed that a population decline could nevertheless be compensated by natural selection acting efficiently over standing variation or by migration of adapted individuals from populations at the margins of the species’ distribution. These findings highlight the importance of within-species genetic heterogeneity in facilitating an evolutionary response to a changing climate.

**One-sentence summary:** “Future genetic changes in *A. thaliana* populations can be forecast by combining climate change models with genomic predictions based on experimental phenotypic data.”

---

Ongoing climate change has already shifted latitudinal and altitudinal distributions of many plant species (1). Future changes in distributions by local extinctions and migrations are most commonly inferred from niche models that are based on current climate across species ranges (2, 3). Such approaches, however, ignore that an adaptive response can occur also *in situ* if there is sufficient variation in genes responsible for local adaptation (4–6). The plant *Arabidopsis thaliana* is found under a wide range of contrasting climates, making it distinctively suited to study evolutionary adaptation to a changing climate (7–9). For the next 50 to 100 years, it is predicted that extreme drought events, potentially one of the strongest climate change-related selective pressures (10), will become pervasive across the Eurasian range of *A. thaliana* (2, 11). An attractive hypothesis is that populations from the southern edge of the species’ range (12) provide a reservoir of genetic variants that can make individuals resistant to future, more extreme, climate conditions (12, 13). To investigate the potential of *A. thaliana* to adapt to extreme drought events, we first linked genetic variation to survival under an experimental extreme drought treatment (14–16). By combining genome-wide association (GWA) techniques that capture signals of local and/or polygenic adaptation (17, 18) with environmental niche models (8, 19), we then predict genetic changes of populations under future climate change scenarios.

We began by exposing a high-quality subset of 211 geo-referenced natural inbred *A. thaliana* accessions (18) to an experimental extreme drought event during the vegetative phase, which killed the plants before they could reproduce (Table S1). After two weeks of normal growth, plants were challenged by a terminal severe drought for over six weeks and imaged every 2-4 days (Fig. 1A) (see Supplementary Online Materials [SOM]). A polynomial linear mixed model was fit to the time-series data to quantify the rate of leaf decay (Fig. 1B-D, Video S1). The genotype deviations from the mean quadratic-term in the model provided the best estimate of this survivorship trait in late stages (Fig. S3, see details in SOM), ranging from −5 to +5 × 10^−4^ green pixels/day^2^. The most sensitive plants survived only about 32 days, while the most resilient plants survived about 15 days longer.

**Figure 1.**
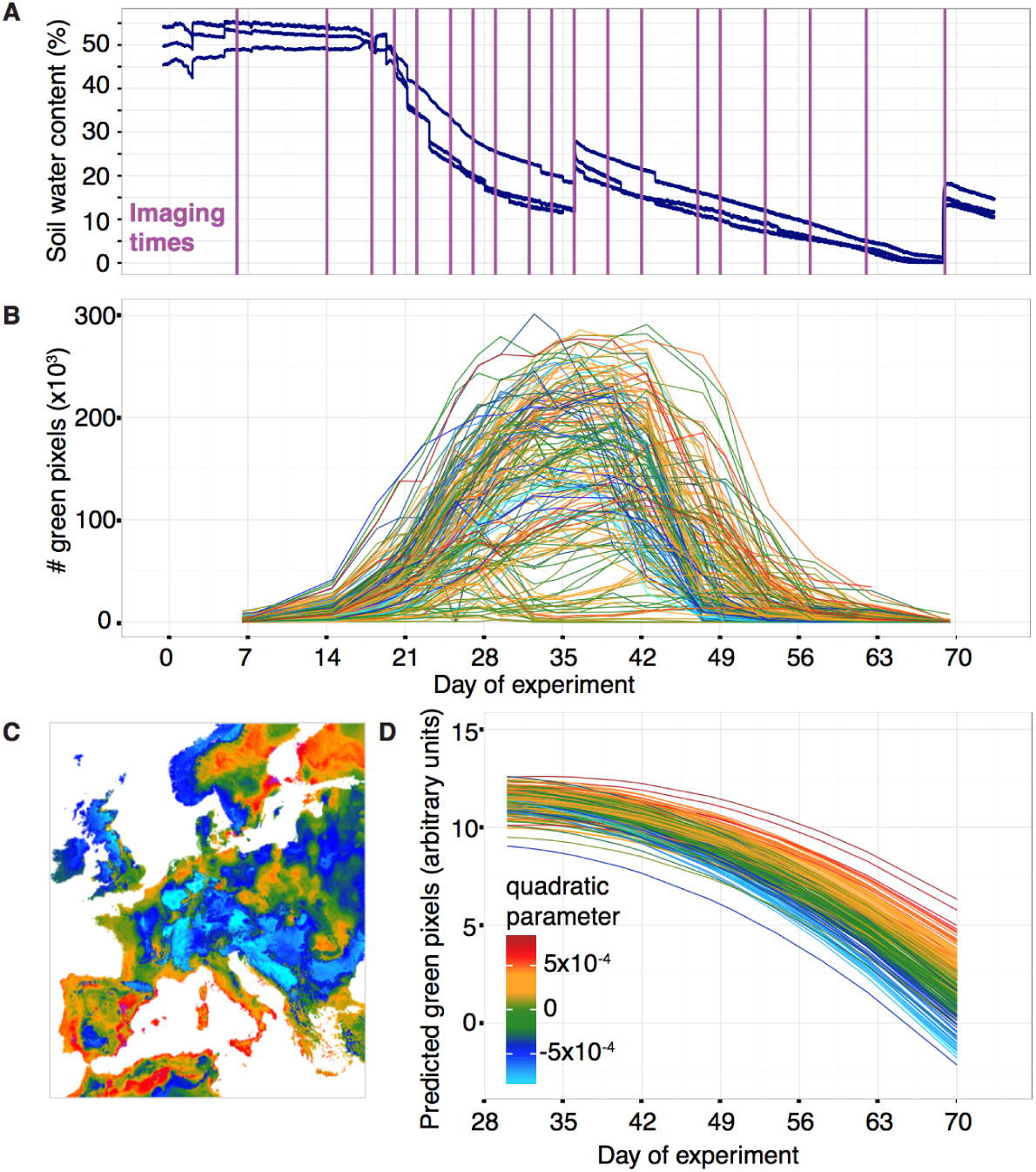
Terminal drought treatment and phenotyping of 211 accessions. **(A)** Soil water content from three sensors placed in three experimental trays regularly distributed in the greenhouse. Purple lines indicate dates of image acquisition. **(B)** Trajectories of total rosette area of 200 randomly chosen pots (see Video S1). Color index according to quadratic parameter in (D). **(C)** Map projection of the environmental niche model prediction of the quadratic parameter (the drought-survival index) in (D). **(D)** Decay trajectory modeled with a polynomial regression, with genotypes as random factors, from the average maximum day of green pixels until the end of the experiment. Each line corresponds to one genotype.

The amount of water available during the drought experiment translates to only about 30-40 mm of monthly rainfall, and as expected, accessions with higher survival come from regions with low precipitation during the warmest season (correlation with climate variable bio18 [www.worldclim.org. ref. (20)]: Pearson correlation, r=−0.19, p=0.005), and specifically with low precipitation during May and June (r≤−0.19, p≤0.005) (see Fig. 2A) (21). To further exploit current climatic data, we used 19 bioclimatic variables and random forest models (22) for environmental niche modeling (ENM) to predict the geographic distribution of the drought-survival index across Europe (Fig. 1C). Surprisingly, we found that individuals with higher drought survival were not only from the Mediterranean, but also from the opposite end of the species’ range in Sweden (Fig. 1C, ENM cross-validation accuracy=89%, Table S10) (21). In contrast to the warm-dry Mediterranean climate, Scandinavian dry periods occur on average at freezing temperatures (Fig. S12). Consequently, precipitation might occur as snow, which is not accessible for plants and produces a physiological drought response (23).

**Figure 2.**
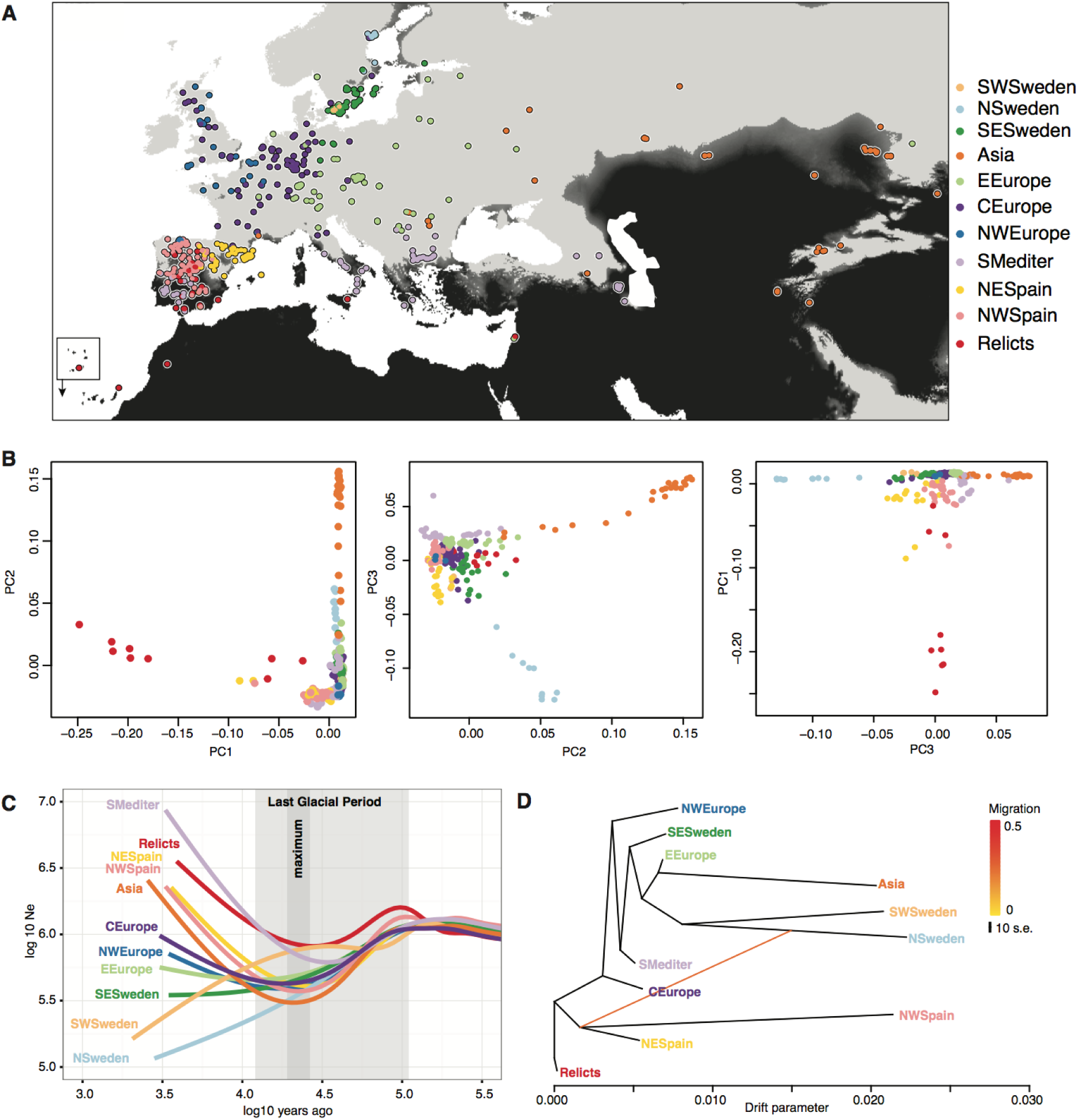
Population structure and history of 762 high-quality genomes. **(A)** Geographic locations and 11 genetic clusters estimated by ADMIXTURE (K11 showed the lowest cross-validation error). Black indicates less than 40 mm of June rainfall (1960 to 1990 average), which corresponds to the amount of water provided in our drought experiment (Fig. 1). Note the presence of black areas in the Mediterranean basin and along the coast in Scandinavia (partially obscured by colored circles). Cape Verde Islands are shown as inset. **(B)** Principal Component Analysis of genome-wide SNPs. **(C)** Effective population sizes in time estimated from MSMC. **(D)** Population ancestral graph and the first migration trajectory using Treemix.

We then studied whether the different populations of *A. thaliana* are locally adapted (5) to low precipitation regimes via an increased drought-survival. Using an extended panel of 762 *A. thaliana* accessions (Table S1) we carried out genetic clustering (24) and studied population trajectories (25) (Fig. 2). This corroborated the existence of a so-called ‘relict’ group (12) and ten other derived groups of relict (e.g. Spanish groups) or other (e.g. Central Europe) origin; likely of the result of complex migration and admixture processes (26). A generalized linear model indicated that genetic group membership explains a significant amount of drought-survival variance (GLM: R^2^=12.8%; p= 4×10^−5^), with the North (N) Swedish and Northeastern (NE) Spanish groups each having on average higher survival than the other groups (t-test p≤0.01). A population graph estimated by Treemix (27) suggested a gene flow edge between the Mediterranean and Scandinavian drought-resistant genetic groups, potentially indicative of historical sharing of drought survival alleles (Fig. 2D). Finally, running an ENM of the genetic group membership with climatic variables from the origin of plants confirmed that the most important predictive variable is precipitation during the warmest quarter (bio18), followed by mean temperature of the driest quarter (bio9), and minimum temperature of the coldest month (bio6) (ENM accuracy > 95%. Fig. S8B and Table S10). As our results indicate that the deepest genetic structure parallels the local precipitation regimes and the ability of populations to survive drought, we expect that areas with the strongest decline in rainfall will see the most turnover in genetic diversity (see Fig.12 Fig. S8) (11).

Because the potential of populations to adapt to drought will depend on the genetic architecture of the selected trait, we identified drought-associated loci with EMMAX (28), a genome-wide association (GWA) method. Although genotype-associated variance (28) h^2^ was 50%, no individual SNP was significantly associated with drought survival (minimum p ~10^−7^, after FDR or Bonferroni corrections p > 0.05) (Fig. S5, Table S3). Significant associations in multiple phenotypes have been detected in similarly powered *A. thaliana* experiments (29). While multiple testing adjustment can over-correct p-values and obscure true associations, the absence of significant associations may also be due to (i) polygenic trait architecture, with many small-effect loci (30) and/or (ii) confounding by strong population structure, consistent with the association of drought survival with genetic group membership.

To test for polygenic adaptation, we repeated the GWA analyses with a model that specifically handles both oligo- and polygenic architectures, BSLMM (31). This model estimates, among other parameters, the probability that each SNP comes from a group of major-effect loci. Around half of the top non-significant EMMAX SNPs were found to have over 99% probability of belonging to such a major-effect group (Fisher’s exact test of overlap, p=3 × 10^−7^; see SOM). We further tested the polygenic hypothesis using the population genetic approach of Berg & Coop (32). The test is based on the principle that if populations diverge in drought-survival due to many loci, there should be an orchestrated shift in their allele frequency. After testing some 60 groups of EMMAX SNP hits of variable size and at different ranks, we detected the most significant signal of polygenic adaptation with the group that included the 151 top SNPs (Table S9). The signal was lost for ranks below the top 300-400 EMMAX SNPs (Table S9). We then compared summary statistics of the top 151 SNPs with background SNPs matched in frequency to avoid GWA discovery biases. The top 151 SNPs showed high F_st_ values, consistent with allele frequency differentiation between populations (Fig. S5). Tajima’s D values were positive (U Mann-Whitney p-value < 0.05), indicating intermediate allele frequencies at the GWA loci (Fig. S5), which could be a result of selection favoring alternative alleles in different ecological niches of the species (33). The top SNPs did not show any evidence for precipitous reductions of haplotypic diversity, as would be expected for hard selective sweeps (34) (Fig. S5). Together these patterns fit the expectations of local adaptation from a polygenic trait controlled by some hundreds loci (35) — theoretically expected to enable a fast response to a new environmental shift (36)

During local adaptation, the relevant loci diverge due to natural selection across populations, which generates a statistical correlation with population groups (37). In this situation, the default correction of population structure applied in GWA might obscure some of the true associations. While F_st_ scans can be useful to identify overly divergent loci across populations, elevated genome-wide F_st_ due to strong population structure can difficult outlier detection (37). as it is in our case (Fig. S4). In order to recover relevant variants that are deeply divergent across populations, we can study the ancestry of each SNP. Using ChomoPainter (38), which relies on linkage disequilibrium information, we segment each genome in question into its different population ancestries (here 11 groups). The first outcome of this analysis was that individuals from NW and NE Spain and, to lesser extent, the Southern Mediterranean (Fig. 2A), have inherited many DNA segments from relict individuals (Fig. S7). Then, in a generalized linear model framework, we test whether the ancestries of individuals at a SNP coincide with the observed phenotypic differences in drought-survival. Performing this “ancestry” genome-wide association (aGWA) and using a permutation correction of p-values (see SOM), we detected 8 distinct peaks (p<0.001, fig. 3A) including over 1,000 significant SNPs (70 SNPs after linkage disequilibrium pruning) (Table S4). The most prominent peak was located on chromosome 5 and explained over 20% of the variance in drought survival (Table S4). There was no overlap in top SNPs between GWA and aGWA because they search for different association signals. Our aGWA resembles other admixture mapping techniques (39). and might be most useful for associations in scenarios of adaptive introgression and local adaptation. To understand the origin of aGWA-identified SNPs, we constructed trees for all concatenated aGWA SNPs and for genome-wide background SNPs. Although the individuals from both the warm (Iberia and relicts) and cold (Scandinavia) edges of the species distribution are far apart in genome-wide SNPs, they are closely related in drought-associated SNPs (Fig. 3B). Overall, this is consistent with a common Mediterranean origin of drought-adaptive genetic variants of both Northern and Southern individuals (Fig. 2D, Fig. 3B), and highlights the relevance of populations at the latitudinal extremes of the species range as a possible genetic reservoir for future climate change adaptation (12).

**Figure 3.**
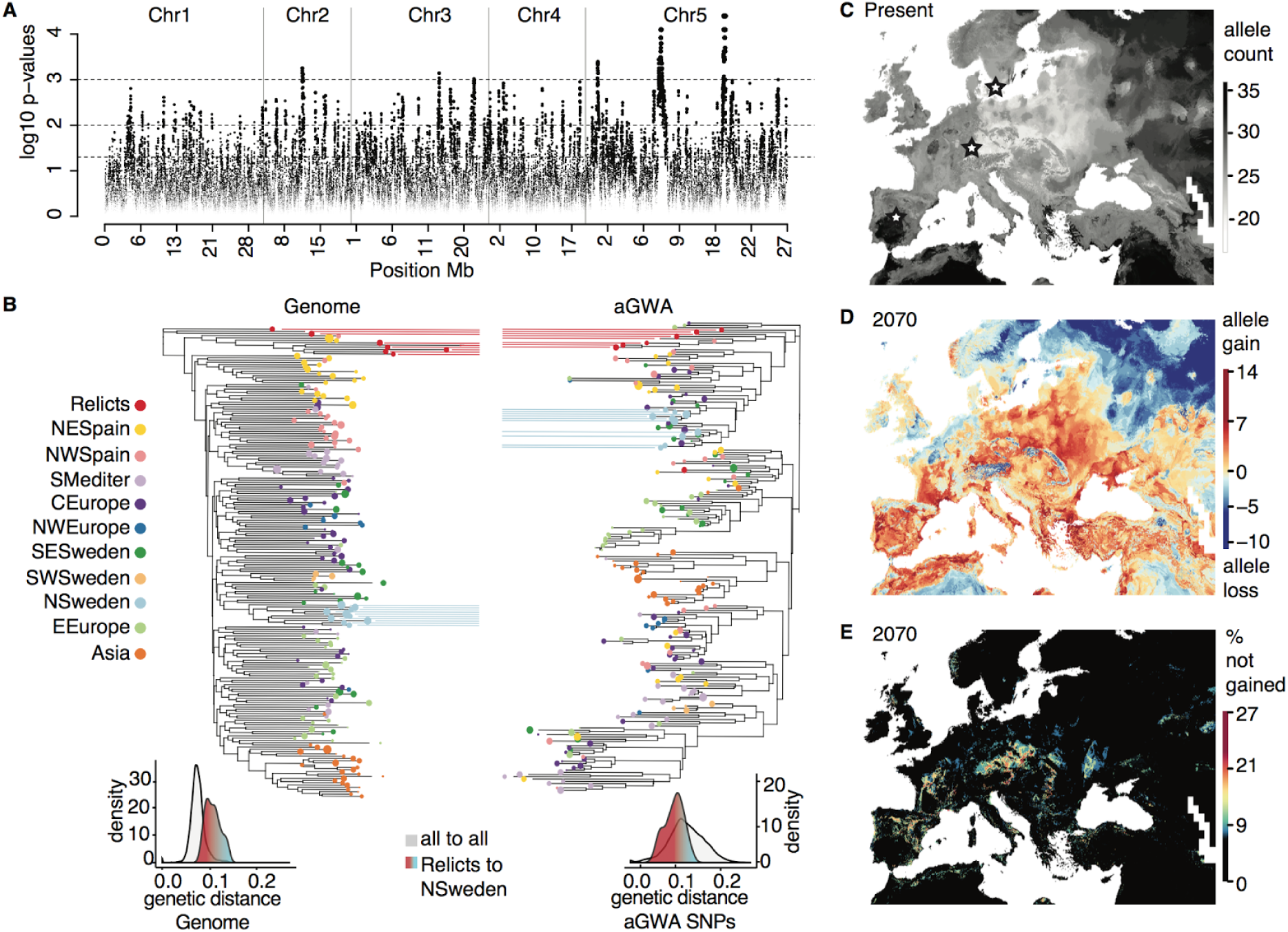
Ancestry GWA of drought survival and environmental predictions. **(A)** Manhattan plot of SNPs from ancestry GWA (aGWA) after permutation correction of p-values. Dashed lines indicate significant thresholds at 0.05, 0.01, and 0.001. **(B)** Top, neighbour Joining phylogeny of 1,000 concatenated genome-wide SNPs compared with a phylogeny of all significant aGWA SNPs (ca 1,000). Colors indicate population clusters (Fig. 2). Relicts and N. Swedish groups are highlighted. Bottom, genetic distances for all or aGWA SNPs. **(C)** Environmental niche models of 70 top aGWA SNPs (after LD pruning), trained with averages from 1960-1990, and then **(D)** used to forecast gain or loss of alleles in 2070 under free migration. (E) The bottom indicates the discrepancy of gained alleles between the geographically constrained (PCA control) model relative to the free migration model.

Depending on the nature of the stress, different mechanisms for drought adaptation can be most advantageous (23, 40, 41). Annual plants, including *A. thaliana*, typically adapt to water stress deficit by accelerating the transition from germination to flowering (escape strategy) (14–16, 41) instead of increasing water use efficiency (avoidance strategy). Previous drought experiments with *A. thaliana* showed variation in both strategies but concluded it predominantly utilizes the drought escape strategy. Our extreme drought experiment focused in characterising the avoidance strategy by means of the drought-survival index, which was linearly associated to precipitation regimes (Fig. S11, Table S6). This trait was not correlated with flowering time of the accessions in unstressed conditions (Pearson correlation, r=0.07, p=0.12). However, we found a positive correlation between drought-survival and flowering time GWA summary statistics of the top 151 SNPs (Pearson correlation, r=0.51, p=1 × 10^−11^, see SOM) — suggesting a weak genetic trade-off (16). Interestingly, we did not find any associated between GWA or aGWA top SNPs and known flowering time QTLs (14–16). but rather a weak enrichment with membrane transporters (see SOM). Adjustment of osmotic balance through cell membrane transport is a drought avoidance mechanism (42) that might also confer cross-tolerance to other abiotic stresses (43), therefore it might be of relevance for Scandinavian *A. thaliana* accessions or other populations in the niche extremes (Fig. S12) (21).

Increased survival to extreme abiotic stresses should confer an evolutionary advantage given the predicted increase in drought frequency and intensity both around the Mediterranean and in Europe, which will constitute a critical hazard for many plants, including *A. thaliana* (2,11). Environmental niche models (ENM), which have been developed to relate species distributions to climate variables, can be used to predict future changes to species’ ranges (2, 3). Ignoring adaptation from standing variation (44–46), however, could lead to overestimates of extinction rates (47–49). By fitting ENM of current climate with SNP data (19). using a similar rationale as in Hancock and colleagues’ “climate GWA” (7), we can predict the most likely genetic makeup under current and future climate conditions. Using such an approach, we trained ENMs with 762 accessions and produced maps of the present distributions of the 151 GWA and 70 aGWA drought-associated SNPs (all ENM 5CV accuracy >92%; Table S3-4, Fig. S13–16). Concatenating the 221 maps, we inferred the most likely individual genotype at each location. At present, individuals from both Northern and Southern edges of the distribution are predicted to harbor more drought-survival alleles than those located in between (Fig. 3C, Fig. S15–16, with the quadratic term in a regression of allele count on latitude being positive at p=10^−3^), corroborating our previous observations. Then, using the trained ENM, we forecast the distribution of the 221 drought-survival alleles in 2070 (rpc 8.5, IPCC, www.ipcc.ch. ref. (20)). While it was expected that populations in the Mediterranean Basin would need to become more drought resistant (11), we predicted a more robust increase in the total number of drought-survival alleles for Central Europe (Fig. 3, Fig. S14–15). This is because rainfall in Central Europe will likely become more similar to that in the Mediterranean by 2070 (2, 11) (Fig. S12).

Because some of the drought-survival alleles are currently not yet present in Central Europe, we speculated that gene migration might be necessary to facilitate adaptation to future conditions (50), An underlying assumption of the ENM is that allele presence only depends on environmental variables, but this assumption, “universal migration”, may not be realistic for future predictions if present distributions are geographically narrow. We therefore included two geographic boundary conditions in the ENM to generate two models that were either more or less “migration-limited” (see SOM). After fitting all possible models and predicting allele distributions with future climate, we calculated the difference of predicted presence per map grid cell between the naïve, free migration ENM and the two geographically constrained ones (Fig. 3D-E). If an allele has currently a narrow distribution or is specific to a certain genetic background, its future presence in an area might not be predicted by the constrained models, even though the climate variables coincide with the SNP’s environmental range. Such a scenario seems to apply to Central Europe, as the deficit in drought-survival alleles predicted by the free over the constrained models was 8-30% (18-66 out of 221) (Fig. 3E; with the quadratic term in a regression of the allele count difference on latitude being negative at p < 10^−10^). Central European populations may therefore be under threat of lagging adaptation by the end of the 21^st^ century.

In the end, for a population to persist, not only the number of drought-survival alleles has to increase, but it has to do so in actual individuals (51). The chance of this occurring will depend on local allele frequencies and the natural selection favouring the drought-survival alleles. Therefore, we studied current allele frequencies at three representative locations with the highest sampling density in our dataset (40 samples within a 50 kilometer area): Madrid (Spain), Tübingen (Germany) and Malmö (Sweden), which are at the southern edge, center and northern edge of the range, respectively. Based on ENM predictions, we calculated allele frequencies from present to 2070. Frequencies are predicted to increase significantly only in the Tübingen population (Student’s t test, p<10^−16^, Table S11), but not in Madrid and Malmö, indicating that these two populations might be already adapted to the future local climate. Because the Tübingen population already has most drought-associated alleles (53% of 70 aGWA SNPs and 90% of 151 GWA SNPs), increasing the number of total favorable alleles in individual genotypes should be feasible, especially since there are single genotypes that have 63% (aGWA) and 90% (GWA) of those alleles already present (see SOM). Starting 50-generations simulations at the present Tübingen frequency of independent drought-survival alleles and assuming a range of selection coefficients, we estimated that a 1-3% of fitness advantage on average would be necessary to increase frequencies to match those of the adapted Madrid and Malmö populations (Fig. S17, see SOM). Such selection could take place efficiently in large populations like the ones of a highly-reproductive weed (51, 52).

Leveraging the model organism *A. thaliana*, we have begun to address key questions to understand the burning issue of climate change effects on biodiversity. We provide evidence for the possibility of adaptive genetic variation to extreme drought events. Harnessing the power of methods that allow polygenic genetic architecture and testing evolutionary hypotheses of natural selection, we detected that relevant genetic variants had been under polygenic local adaptation and were more abundant at the edges of the species range. Extreme adaptation at range edges might indeed be critical for a species’ persistence under climate change. Although many aspects of future adaptation are not considered here, namely non-drought related or seasonal climate change (51). biotic interactions, phenotypic plasticity, or novel adaptive mutations (53), our spatially explicit analyses emphasize the potential of adaptive evolution from standing variation to ameliorate climate change’s detrimental effects.

## SUPPLEMENTAL MATERIALS

Methods and any associated references are available in the online version of the paper. [link]

Supplementary text

Figs. S1 to S17

Video S1

Tables S1 to S1

References (* to *)

### URLs

Code for image analysis pipeline available at http://github.com/MoisesExpositoAlonso/hippo. Code for ancestryGWA available at http://github.com/MoisesExpositoAlonso/aGWA. Bioclimatic data used in the paper accessible through an R package stored in http://github.com/MoisesExpositoAlonso/rbioclim. Phenotypic datasets available at dryad [*link here*]. Genomes are available at http://1001genomes.org/data/GMI-MPI/releases/v3.1/.

## Acknowledgements

We thank I. Henderson for the recombination map, R. Wedegärtner for assistance with the greenhouse drought experiment, the Petrov, Coop, Ross-Ibarra, Gaut and Schmitt labs for discussions. We thank J. Lasky, X. Picó, A. Hancock, H. Thomassen, T. Mitchell-Olds, J. Mujica, P. Lang, and D. Seymour for comments and the Weigel and Burbano labs for discussion. This work was supported by ERC Advanced Grant IMMUNEMESIS to DW and the President’s Fund of the Max Planck Society, project “Darwin” to HAB.

## Author contribution

MEA conceived and designed the project. GW and FV helped and advised on image phenotyping and FV provided additional phenotypes. MEA and WD performed chromosome painter analyses. MEA performed the drought experiment, processed the image data, and designed and carried out the statistical analyses. DW and HAB advised and oversaw the project. MEA wrote the first draft and together with HAB and DW wrote the final manuscript with input from all authors.

## SUPPLEMENTARY TEXT

### 1. Experimental design and biological material

#### 1.1 Choice of accessions from the 1001 Genomes resource

The 1001 Genomes project has released resequencing data for 1,135 natural inbred lines, also called accessions (http://1001genomes.org). We applied several filters to select the most informative, least biased accessions for our experiment, (i) The first filter removed 176 accessions with low quality genome information, < 10X genome coverage and < 90% congruence of SNPs called from Max Planck Institute and Gregor Mendel Institute pipelines (1). (ii) The second filter removed 244 nearly-identical accessions, many from N. America. For this, we calculated pairwise genome-wide identity-by-state differences using PLINK v1.9 (2). When pairs differed in less than < 0. 01 changes per polymorphic site, we randomly removed one member of the pair. The overlap between (i) and (ii) was 762 accessions (Fig. S1, S2, Table S1). For geographic analyses in the native range (e.g. environmental niche models), we used the 729 accessions that were within 50°W to 100°E longitude, i.e. Eurasia (see section 4.2.1). For the terminal drought experiment, we used 211 of these 729 accessions. The seeds were progeny of the 1001 Genomes collection seed stocks obtained from the ABRC Stock Center (CS78942).

#### 1.2 Greenhouse terminal drought experiment

The 211 accessions included both vernalization-requiring, slow-flowering and vernalization-independent, fast-flowering ones. Because the phenotype of interest in our study was the variance in survival during vegetative stages under terminal drought conditions, we applied a terminal severe drought without prior vernalization. Because of the difficulties associated with disentangling drought-induced mortality and reproduction-associated senescence at the end of the plant life cycle, our study focused on drought stress during the vegetative stage. (Note that onset of flowering, or flowering time, was not a confounding factor. See section 2.2).

Seeds were aliquoted in Eppendorf tubes, suspended in 1% agar solution, stratified in a 4°C dark room for 5 days to promote germination, and then pipetted into pots filled with sieved soil (CL-P, Einheitserde Werkverband e.V., Deutschland). When multiple seeds germinated per pot, all but one were removed at random. We sowed 8 replicates per genotype in 49 trays of 8x5 cells (5.5 × 5.5 × 10 cm) using a randomized incomplete block design. We excluded corner cells, where edge effects are strongest.

During the first two weeks after sowing (defined as day 0), trays were watered close to soil saturation once every 3 days, with temperature maxima from 20 to 25°C under16 hours natural and supplemental light. After this period, seedlings were challenged with a terminal drought, with “recovery waterings” after 3 and 6 weeks, in order to increase the variance in survival. The overall watering during the drought period (4 l in each tray of 40 × 60 cm), corresponded to approximately 33 mm of rainfall (4,000 + 4,000 cm^3^ water/ 2400 cm^2^ surface = 3.3 cm). We monitored water content using moisture sensors (Parrot SA, Paris, France) (see water content graph in Fig. 1A). We monitored rosette green area by imaging at 20 time points (Fig. 1A) using a customized system (see below).

#### 1.3 Experiment under optimal growing conditions

In a first experiment, we grew the same 211 genotypes under optimal watering and nutrient conditions and monitored vegetative growth by image analysis (Vasseur et al. submitted [*link to preprint or in press article will be added here*]) (see Table S5 for a description of the 24 traits extracted from the images). This set of traits was used to investigate whether variation of drought-survival index was correlated with growth under optimal conditions.

### 2. Drought phenotyping

#### 2.1 Image analysis pipeline

Images were taken using a Panasonic DMC-TZ61 digital camera and a customized closed black box at a distance of 40 cm from the tray. This produced very consistent images in terms of illumination (only from in-camera flash) and focal distance to the plants.

We extracted leaf area per plant over time using the imaging module Open Computer Vision in Python (3) (Video S1), with these steps: (i) 5 pixel mean denoise of the whole-tray image, (ii) Fixed Hue Saturation Value (HSV) segmentation of “green” values. The threshold values were determined heuristically by comparing the HSV ranges of leaves at five timepoints (i.e. early and mature), (iii) Cropping of each pot to extract individual plant images, (iv) Counting of green pixels (for more specific details and code visit http://github.com/MoisesExpositoAlonso/hippo). Pots with green pixels but without plants were excluded after careful visual inspection of all images.

#### 2.2 Drought index

After determining the peak of green area for the majority of pots, we modeled the daily number of green pixels per pot. Several different models, including up to third order polynomial models, several error correction factors, either raw or genotype averages, were tested. All models were ranked based on parameter convergence in an MCMC walk and AIC values. The final chosen model was a generalized linear mixed model with Poisson link of the form:

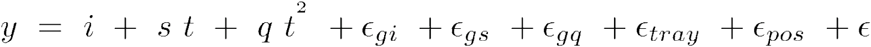

where green area, *y*, was the response variable, and an intercept (*i*), slope (*s*), and quadratic coefficients (*q*) with time (*t*) were fitted as fixed effects. Genotypes were treated as random factors, that is allowed to deviate from the main trends, following a normal distribution (0, lσ_g_). Tray block and position within the tray grid were fitted as random factors following also a normal distribution. To estimate these parameters, we performed 10,000 iterations in a Monte Carlo Markov Chain (MCMC) and 1,000 burn-in using the glmmMCMC R package (4).

The variance from all genotype-dependent components relative to the total phenotypic variance was ~10%. This might be an underestimate since certain variance is due to inaccurate phenotyping: either optical and illumination distortion, or overlap of neighboring plants. Genotype values of the three parameters of interest (intercept, slope, and quadratic coefficient) were used for Genome Wide Association (GWA) and downstream analyses. Additive genetic variance was estimated from linear mixed models using a kinship matrix (see GWA section (3.3)). The intercept, slope and quadratic deviations had narrow-sense heritabilities (h^2^, or kinship-associated variance) of 0%, 0%, and 50%; respectively. We chose the latter as the drought-survival index. This parameter informed about survival during the late stage of the experiment, as can be observed from a high correlation between the drought-survival index and the the raw green pixels in the final monitored days (Fig. S3).

Because the drought-survival index could depend on the developmental stage of the plants when the drought treatment started, we computed the pixel decay polynomial model with and without a covariate of flowering time under optimal conditions (indicative of developmental speed; see source of the phenotype in section 1.3 and phenotypic correlations in section 3.4.1). The small effect of flowering time in the model (0.01% of the variance) confirmed that the measurements were not biased, therefore we removed flowering time from the final model.

In order to provide an intuitive understanding of the drought survival index, we looked at the relationship between the index and the last day on which a plant was clearly alive, defined as the last day with at least 5,000 green pixels left. The relationship between the drought-survival index and the last living day was highly significant (p < 10^−16^). The most sensitive plants survived for 32 days, and the most resistant were alive for 15 days longer.

### 3. Quantitative genetics analyses

#### 3.1 Population structure

From the vcf file containing SNP calls from the 1001 Genomes project (http://1001genomes.Org/data/GMI-MPI/releases/v3.1/), we identified SNPs with a genotype calling rate >95%, which resulted in ~4M SNPs in 762 accessions (section 1.1). We defined genetic clusters with ADMIXTURE v1.2 (5) (Table S2). As a model-free alternativeto ADMIXTURE, we used PCA implemented in PUNK v1.9 (2). The first three axes explained 16.0%, 9.6% and 7.9% of the total genetic variance. ADMIXTURE clustering and PCA were used to understand population structure and to relate it to phenotype variables. We assessed population splits and migrations with a population ancestral graph using TreeMix v. 1.12 (6), a tree based on genome-wide allele frequency differences across populations. Additionally, we calculated a proxy of local genetic diversity (7) per location using the number of polymorphic sites in the geographically closest genome.

##### 3.1.1 Association of genetic group membership with drought

Using the ADMIXTURE membership probabilities of each genome, we carried out univariate linear regressions with the drought survival phenotype. The groups that yielded positive relationships were NE Spanish (p<0.05), Mediterranean (p=0.06), and the N. Swedish groups (p<0.001). The groups negatively associated were Central Europe (p=0.06), Asia (p<0.001), and E. Europe (p<0.001). This broadly coincided with the map of drought-survival prediction (Fig. 1D, S11). We found that only PC3 was significantly associated with the drought survival phenotype (GLM R^2^=0.076; p=5.15×10^−5^). The N. Swedish and NE Spanish groups showed particularly low values in PC3 than the rest (Fig. S8).

#### 3.2 Coalescent rates over time

Only the accessions with ≥90% of membership probability in one of the genetic groups were used. Using MSMC software (8) (http://github.com/stschiff/msmc), we performed within- and cross-genetic group coalescent analyses by contrasting two pairs from different genetic groups. In total 333 analyses were performed, with each genetic group being tested at least 3 times. The results were summarized using a smoothed generalized additive model in R (Fig. 2C).

#### 3.3 Genome wide associations (GWA)

##### 3.3.1 Linear Mixed Models (LMMs) with EMMAX

We used 879,654 biallelic SNPs with a minimum allele frequency (MAF) of 5% for genome wide association (GWA) using EMMAX (9). We carried out GWA for all climatic variables and 11 phenotypes (Table S5). The GWA is based on linear mixed models that test, one by one, each of the SNPs, and correct the results by population structure using a random factor with a variance/covariance kinship matrix built from genome-wide SNPs. This is an appropriate method to correct for coancestry in *Arabidopsis thaliana* (10).

To rule out the possibility that drought survival measurements were dependent on the developmental stage of the plant during the experiment, we carried out the GWA with and without a covariate of flowering time that had been scored in controlled conditions (proxy of developmental speed; section 1.3). The top SNP hits were the same with or without this covariate, and we only show results without the covariate. To account for familywise error in GWA we used Bonferroni correction (p value × number of SNPs) and the Benjamin-Hochberg false discovery rate correction (11). The kinship-associated variance of drought-survival — an approximation of narrow sense heritability, h^2^, was 49%. When we fit a kinship calculated from only the 151 top polygenic GWA SNPs (see section 3.5.2), the estimate of h^2^ was 52%. This is probably a better estimate than that from the genome-wide-based kinship matrix, as the putatively causal SNPs are better “tagged” in the 151 SNPs kinship matrix.

After calculating genome-wide F_st_ (12) and Tajima’s D (13) with PLINK v1.9 (2) and likelihood of a selective sweep with SweeD (14), we investigated the enrichment of the top SNPs in the upper tail of the distributions of those statistics (Fig. S4) (Table S3, rank columns).

##### 3.3.2 Bayesian Sparse Linear Mixed Models (BSMLMMs) with GEMMA

The Bayesian Sparse Linear Mixed model (BSLMM) implemented in GEMMA (15) accommodates both poly- and oligogenic architectures in a GWA framework. It models two effect hyperparameters, a basal effect, *alpha*, that captures the fact that many SNPs contribute to the phenotype, and an extra effect, *beta*, that captures the stronger effect of only a subset of SNPs. The parameter measuring the probability of having another extra effect, *gamma*, can be used to prioritize SNPs (personal instructions from the author X. Zhou). Over 40% of the top 151 SNPs from EMMAX were found to have over 99% percentile of the gamma inclusion probability in GEMMA (Fisher’s exact test odds ratio = 17.21, p=3×10−^7^). The estimate of realized heritability with BSLMM was 50%, which is in agreement with the EMMAX analyses. The 95% highest posterior density (95%HPD) from 1,000 MCMC steps ranged from 25-85%.

#### 3.4 Multivariate analyses of phenotypes and GWA summary statistics

For a description and sources of all variables used, see Table S5.

##### 3.4.1 All pairwise correlations

We computed all-against-all Pearson product-moment correlation coefficients among accession line means (n= 211 accessions) of phenotypic and climate variables (Table S5, S6). To study genetic correlations, we performed the same analyses with SNP effect sizes (n= 151 drought-associated SNPs) estimated from multiple GWA (Table S7).

The phenotype correlations (Table S6) showed that the drought-survival index was negatively correlated with reproductive allocation and number of seeds (r < −0.16, p<0.02), suggesting a fitness trade-off between stressful and optimal growth environments. Drought-survival was not correlated with flowering time (r=0.07, p=0.12) nor plant size (rosette area and dry mass, r<0.12, p>0.07).

Drought-survival SNP effects negatively correlated with the SNP effect sizes of most precipitation variables separately (r < −0.4; p < 10^−8^, Table S7), indicating that alleles that increased drought survival were found in more arid geographic regions, i.e. regions with high temperatures and lower precipitation at different times of the year. Drought-survival SNP effects were also positively correlated with SNP effects of rosette area, dry mass, and flowering time (Table S7). These analyses have two-fold interests: (1) GWA-estimated effect have been corrected by population structure, thus correlations should not reflect phenotypic differences caused by drift of populations. (2) SNPs can have pleiotropic effects and this can limit adaptation due to genetic constraints (see section 4.2.4.3) (16).

##### 3.4.2 Canonical Correlation Analysis (CCA)

We further utilized Canonical Correlation Analysis (CCA) to decompose environment-phenotype associations of SNP effects. This was done for all genome-wide SNPs (n=~800,000) and for the 151 drought-associated SNPs (Table S8).

CCA of genome-wide SNPs revealed the first canonical correlation axis (CC1) to be driven by lower flowering time (T_repro, loading=−0.77), lower rosette dry mass (loading=−0.76) and higher annual temperature (bio1, loading=0.5). CC2 indicates that lower plant photosystem stress (FvFm, loading=0.60) is related to higher mean temperature of the wettest quarter and higher precipitation seasonality (bio8, bio15, loadings>0.25). CC3 shows that lower drought survival (loading=−0.58) effects are related to higher precipitation in the driest (bio17, loading=0.44) and warmest quarters (bio18, loading=0.35).

CCA of 151 top GWA SNPs yielded a first canonical correlation coefficient of 0.99, with a phenotype canonical variate driven by lower drought survival, higher rosette area and dry mass (loadings >0.75), and a climatic canonical variate dominated by higher precipitation during the wettest month (bio13) and wettest quarter of year (bio16) (loadings >0.75).

#### 3.5 Polygenic adaptation signal

##### 3.5.1 Classic Q_st_-F_st_ comparison

Q_st_/F_st_ ratios have been proposed as an appropriate indicator of local adaptation in *A. thaliana* (17) and related species (18). Genome-wide F_st_ across the eleven population was computed from 211 genomes using vcftools (v0.1.12b) (19). We estimated the mean and confidence intervals based on the standard error of the mean, obtaining a mean F_st_ = 0.042 (95% cumulative distribution = 0.360). We calculated Q_st_ for the drought-survival index as the between-genetic group variance divided by the total variance. We used the MCMCglmm function in R with a 10,000 steps chain, 1,000 steps burn-in, and fitting the genetic group as random effect. This resulted in a Q_st_ = 0.143 (90%HPD = 0.052 - 0.338). When the variance across populations was done using the NE Spanish and the NSweden (population groups hypothetically under local adaptation) against the rest, we obtained Q_st_= 0.377 (0.047 - 0.987). We thus concluded that a significant Q_st>_ > F_st_ signal is only observable at the individual level when the hypothetical populations that underwent local adaptation were oriented in the calculation of the variance.

##### 3.5.2 Berg & Coop methodology

We tested for a polygenic adaptation signature following Berg & Coop (20), an extension of the Q_st_/F_st_ ratio test based on SNP frequency per population and effect sizes as estimated from a GWA analysis. We used different groups and numbers of ranked SNPs after pruning linked SNPs (r^2^ > 0.6), to learn about the robustness of this test and the apparent number of SNPs that contribute to the signal (Table S9). Since this test does not use direct phenotypes but calculates the average phenotype per population based on allele frequencies of GWA SNPs, we could perform the test with 762 high quality accessions. Since results did not vary between 762 and 211 sample analyses, we only report the analyses with the 762 genomes (Table S9).

#### 3.6 ChromoPainter and ancestry GWA

We ran ChromoPainter version 2.0.7 (available at http://paintmychromosomes.com) (21) on the 762 genomes dataset, after imputing missing genotypes with Beagle version 3.3.2 (22) using default parameters.

ChromoPainter analyses require a “training” run to estimate several hyperparameters. We ran 10 expectation maximization iterations on chromosome 2 (the smallest chromosome). We informed ChromoPainter with a published recombination map of *Arabidopsis thaliana* (23) that we reshaped to our SNP dataset. We used the command:

ChromoPainterv2 -i 10 -in -iM -j -g haplotypefile -r recombinationfile -a 0 0 -t labelfile

We used the output hyperparameters to run ChromoPainter on all chromosomes in an unsupervised all-to-all genomes mode, with the command:

ChromoPainterv2 -n 4.737068 -M 0.000421 -j -g haplotypefile -r recombinationfile -a 0 0 -t labelfile

##### 3.6.1 Global proportion of ancestral chromosome segments

To study the ancestry relationships of each of the genetic groups, we counted the number of chromosomal segments (termed “chunks” in the original ChromoPainter paper (21)) that each genome “received” from all other genomes. The segment varied in size depending on local recombination rates and between genomes, but *a posteriori* analyses indicated that the median size was in the order of magnitude of kilo and megabases. To make the counts more informative, we show boxplots per ADMIXTURE group rather than counts per individual (Fig. S7 A-K). This showed, for example, that NW Spain, NE Spain and S. Mediterranean (the latter to a lower degree), were “painted” mostly by relict DNA segments. Next, we tried to infer how well the drought survival of an individual correlated with the number of segments inherited from a certain ancestry. This indicated that only N. Sweden and relicts passed DNA segments that were correlated with the drought-survival index of the receiving individual (Fig. S7L). The Pearson correlation coefficient was calculated excluding the individuals from the same admixture group as the predictor.

##### 3.6.2 aGWA for admixture mapping

If populations are locally adapted, F_st_ outlier scans can be used to identify genetic variants under divergent selection (24, 25). However, when populations get isolated and diverge genetically, as it is the case in *Arabidopsis thaliana*, F_st_ values are shifted to high values across the entire genome even when subsequent admixture happen, making the identification of outliers difficult (Fig. S4) (24). Thus, we must rely on linkage disequilibrium and identity by descent to find DNA segments characteristic of the different populations. If subsequent but incomplete admixture occurred between the locally adapted populations, it is expected that the individuals that retained the DNA segments responsible for local adaptation, would show the largest phenotypic differences. This is the principle of admixture mapping (26).

With the above rationale, we developed an admixture mapping technique (26) repurposing the output of ChromoPainter. The “painted” genome matrix produced by ChromoPainter has 762 states (one per individual in the analysis) and we repainted it into a genome matrix of 11 states (the genetic groups from ADMIXTURE analysis, which are geographically and environmentally separated). We then computed a regression of the drought-survival phenotype on the population group specific to a SNP as:

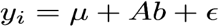

Where *y* is a vector of *i=1…211* individual’s phenotypes, *μ* is the mean phenotype, *B* is the 211 × 11 design sparse matrix of the ancestry states, *b* is a 1 × 11 vector of effects of each ancestry has in the mean phenotype, and *ε* is the uncorrelated random residuals assumed to be normal. This model was repeated for each SNP in our dataset (~2 million imputed and ‘painted’ SNPs, see section 3.6). We report R square of and p-value of each SNP model (Table S4). Since we already knew that the phenotype is associated with the membership assigned per individual, we expected that the membership of any random SNP would be on average also associated, because of linkage resulting from common ancestry. Therefore, we implemented an empirical p-value distribution correction to only detect those SNPs whose ancestry explained an even larger proportion of variance than the whole-genome ancestry. The permutation was done within each individual genome shuffling the SNP states at a distance of 1,000 to 10,000 SNP positions — defined from analysing the typical size of “homogeneously painted DNA segments” (code are available at http://github.com/MoisesExpositoAlonso/aGWA). We permuted the dataset 1,000 times and repeated this “aGWA” analysis to build p-value distributions. Since the nature of the associations is very different from that of a standard GWA analysis, we did not expect and did not find any overlap of top aGWA SNPs with the top SNPs from regular GWA. The closest was a regular GWA SNP that was 8 kb away from an aGWA SNP. The closest gene to both encodes a defensin-like protein; a family of proteins with broad anti-fungal and anti-bacterial activity (27).

Our approach is conceptually related to admixture mapping in humans, which has focused on local enrichment of Neandertal- and Denisovan-like variants, and which has led to the identification of in TLR immunity genes (28) as adaptive. It has also helped to increase the power for detection of background-dependent disease risk in humans with mixed ancestries, e.g. African-American individuals (29), or other more complex mixtures (30). Such approaches constitute a powerful tool for understanding the genetic basis of local adaptation when complex demographic scenarios of admixture exist.

###### 3.6.2.1 Phylogeny of aGWA SNPs

To learn about the distribution and shared ancestries of the drought-related alleles, we computed a neighbour joining phylogeny of all concatenated SNP hits from aGWA (p<0.001) and compared it with a genome background phylogeny of 1,000 randomly chosen SNPs (Fig 3B). This revealed that the N. Swedish groups and Mediterranean relicts were across the genome more distant from each other than the average between-group distance (Student’s t test, p<2×10^−16^) (Fig. 3B), much closer than the average when considering only the aGWA SNPs (Student’s t test, p<2x10^−10^). The same analyses showed also higher affinity of N. Swedish and NE Spanish populations (Student’s t test, p<10^−10^).

#### 3.7 Test for annotation enrichment

Using the TAIR10 gene annotation of *Arabidopsis thaliana* (available at arabidopsis.org/portals/genAnnotation/functional_annotation/), we tested whether there was a specific annotation class enriched in our GWA and aGWA hits. The most suggestive genes overlapping with the 151 top GWA hits were the nitrate transporter gene *NRT1.8*, which among other functions mediates cadmium tolerance and is related to ABA transport (31–33), the *CATION/CARNITINE TRANSPORTER 4* (*OCT4*), which mediates homeostasis of metabolites and promotes lateral root formation (34), and the sugar transporter gene *SWEET8*, which is upregulated during salt stress (35). On the other hand, the strongest peak fell inside the *CATION EXCHANGER 9*, a gene important for homeostasis of K^+^, Na^+^ and Mn^++^ that confers salt tolerance when introduced into yeast (36). An empirical distribution test based on random draws of genes showed, however, only marginal enrichment. The 30 genes defined by the 151 top GWA SNPs were weakly enriched for cell membrane transport (6/30; p=0.01), and the 23 genes defined by the 70 top aGWA SNPs were only very marginally enriched for membrane transport (7/23; p=0.06). Testing for overrepresentation with PANTHER (www.pantherdb.org) and including genes adjacent to the GWA and aGWA SNPs revealed weak enrichment of aGWA genes for ferredoxin metabolic process (p=0.03) and vesicle-mediated transport (p=0.05), and of GWA genes for growth-related functions (p=0.0007) and metabolite biosynthetic processes (p=0.0002). It is difficult to know what to conclude from this, but the most noteworthy finding is probably that there was no link to flowering time, in contrast to previous QTL and GWA studies of *A. thaliana* response to drought (37–39).

### 4. Environmental and forecasting analyses

#### 4.1 Environmental data

The environmental data comprised the Last Glacial Period (LPG, ~22,000 years ago), recent averages from 1960-1990, and two 2070 climate projections of contrasting socio-economic scenarios, the 2.6 and 8.5 CO_2_ representative concentration pathways (rcp) (40, 41). The data was retrieved from www.worldclim.com v.1.4 (ref. (42)). It consists of 19 bioclimatic variables at 2.5 minutes geographic resolution (code to retrieve and process data available at http://github.com/MoisesExpositoAlonso/rbioclim).

#### 4.2 Environmental Niche Models (ENM)

We carried out ENM with a number of response variables (for summary statistics see Table S10), namely the drought-survival phenotype, flowering time, the genomic principal component axes, the discrete population groups, the local genetic diversity, and the SNPs identified in GWA and aGWA analyses.

##### 4.2.1 Geographic areas used and niche limits

To train ENM, we removed accessions from Japan and from N. America, as they are recent introductions (43) and might not reflect long-term climate adaptation. In addition, the sampled locations used to trained the models were within 15 to 63° N and 23°W to 88°E longitude, but we only predicted in a reduced area, from 34 to 63° N and -10.5 to 35°E, to avoid extrapolation of data. Predictions for the last glacial maxima were masked in those areas that were likely tundra or ice sheet at the time (<5°C and <0°C annual temperature, respectively), as they are presumably artifacts.

Because the sampling in the 1001 Genomes project was not even across the species range, predictions for underrepresented regions such as N. Africa, the Middle East, or Russia, must be taken with caution. In order to be explicit about for which areas we could make the most robust predictions, we show the sampling density per 1° × 1° latitude × longitude grid, which varies from around 1 to 60 individuals (Fig. S8D). and plot trends of predicted values against other variables, such as latitude or climate variables (e.g. Fig. S9–S11). only at those locations where there is at least one sample.

Finally, it is worth noting that even for the most pessimistic climate change scenario (rcp 8.5), the values of annual precipitation (bio1) and the precipitation during the warmest season (bio18) were always above the present minimum precipitation values where *A. thaliana* is currently found (see Fig. S12). Therefore, we expect that transgressive phenotypes are not required to survive future climates.

##### 4.2.2 Random Forest models

After trying different methodologies, including generalized linear models, MaxEnt, and linear discriminant models, we opted for random forest models because they are nonparametric, nonlinear, allow both continuous and discrete response variables, and are computationally efficient (44). Additionally, the implementation of an “importance” parameter of each predictor variable available in the *randomForest* R package makes ranking of variables straightforward. To mitigate the overfitting problem typical of machine learning methods, a 5-fold cross validation procedure was used. We randomly divided the dataset in five parts, used four parts as training dataset and one part as testing dataset, and repeated this five times. Reported accuracy from cross validation was the R^2^ of a linear model between observed and predicted values for continuous variables, and the rate of successful assignment of categories relative to the total number of observations for discrete variables. To build the final forest, a total of 50 classification or regression trees per cross-validation set were used, and six variables were tested for each classification split.

##### 4.2.3 ENM of genetic groups

We modeled the presence of population structure as a discrete response variable in ENM; either using eleven genetic groups as states, or the two relict and non-relict states.

In order to formally quantify the relevance of genetic group membership, we calculated the percentage of map grid cells that each genetic group occupies. For this we only considered areas where at least one genome per 1° × 1° latitude × longitude was observed (Fig. S8A) and where tundra or ice sheet are not expected (important for LGM comparisons).

When we used the present-data trained relict/non-relict ENM with past climate data from the last glacial maxima, we found that relicts likely occupied almost a quarter of the non-glaciated areas, compared to less than 2% today (Fig. S8D). in agreement with genomic inferences of higher effective population size in the past (Fig. 2C). The reason that the relicts’ environmental niche is predicted with 100% accuracy under 5-fold cross-validation (5CV) is likely that the local number of relicts individuals is low, 26 accessions out of 762, and because their niche is very restricted.

Under a future high CO_2_ increase socio-economic scenario, the ENM with 11 genetic groups predicts that the S. Mediterranean group will expand most dramatically into Central-European areas, replacing groups currently occupying these areas (Fig. S8 C, F). Although these models are not mechanistic, they illustrate that genetic groups from the Mediterranean and from temperate areas have contrasting environmental niches and thus might replace each other under future climate warming.

##### 4.2.4 Genome Environment Models (GEMs)

All 151 GWA and 70 aGWA SNPs were modeled as a bivariate discrete variable (drought-sensitive and drought-survival allele) in a random forest. The prediction accuracy and the most important predictor for each model below is shown per SNP in Table S3 and S4.

After modeling presence/absence of each drought-survival SNP, we projected the present inferred allele distributions in a map and then summarized all maps by intersecting them. In this way, we generated a continuous map surface of the total number of drought-resistant alleles in a given location. Ancestral GWA and GWA models showed overall similar patterns (Fig. S13–14). but the latter were more biased to Western areas. This is likely due to GWA SNPs suffering from high-frequency bias, making them more likely to be present in geographic areas with more samples (Fig. S8). After we had trained the models with present data, we used them to predict allele distributions in 2070 under low and high CO_2_ increase scenarios. While patterns were similar in both scenarios, for further analyses we used the most “pessimistic” high CO_2_ increase scenario to be able to show main trends more clearly.

###### 4.2.4.1 Migration assumptions

For each SNP we trained three models in order to overcome the “universal (or free) migration” assumption, implicit when using a current climate-trained ENMs with future climate data (e.g. (45)). Although typically the free-migration model may not be entirely appropriate for predictions, it might be more realistic for cosmopolitan species with continental-scale migrations in the recent past, as is the case of *A. thaliana* (43). Nevertheless, we designed two additional models to account for limited migration. The free model includes only the 19 bioclimatic variables as predictors of the drought-survival alleles. The first geographically-controlled model includes in addition the first three PC genomic axes as predictors (Fig. S8G-J). in an attempt to limit prediction allele presence to geographic areas where the genomic background that they reside on is present today. The second geographically-controlled model, which is even more restrictive, includes latitude and longitude together with the 19 bioclimatic variables. For all models we not only show the predicted maps (Fig. S13–14) but also provide residuals of predicted *vs* observed (empirical) number of alleles in the locations where we have a sample. We also show their relationship with latitude (Fig. S15–16).

###### 4.2.4.2 Allele frequency change predicted by GEMs

We took 40 individuals approximately within 50 km of each other at three locations with the highest density of samples in our dataset: Madrid (Spain), Tübingen (Germany) and Malmö (Sweden) (Fig. 3C, Fig. S8). We tested overall future allele frequency changes of all SNPs per population as well as SNP-specific allele frequency changes.

First, we calculated the mean allele frequency differences between future (rcp 8.5, 2070) and present predictions. This proved to be significant in most locations and models (Table S11), although the direction of change was different between the two edge populations, Madrid and Malmö, and the Tübingen population from the center of the range. The former showed a decrease or a steady state in allele frequency, and the latter showed a highly significant increase in all models and SNP subsets (Table S11).

Secondly, we calculated the differences in frequency (F) between present (*pres*.) and future (*2070*) populations per SNP and tested the difference using a Student’s t test and a pooled standard error (*se*) of the frequency measurements:

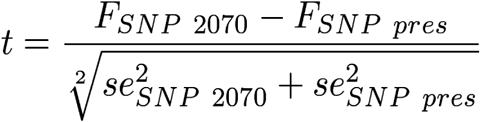

This not only revealed the main frequency change trend, but also the distribution of differences in alleles (see histograms in Fig. S13–14). It corroborated the general trend observed for all SNPs (Table S11) and in addition showed that the global distribution of allele frequency changes in Tübingen is skewed to the right in some SNPs (increase of drought allele frequency).

###### 4.2.4.3 Possible genetic trade offs of drought survival and flowering time

Contrary to our expectations, there were areas in the Mediterranean that were predicted to lose drought-survival alleles under climate change (Fig. S9–10). These are areas that already suffer today from low precipitation (reached the zero in summer, Fig. S12) and will probably not become much drier in summer. On the other hand, temperatures will keep increasing, which likely will demand an acceleration of flowering time (in trade-off with drought avoidance). Predictions at the phenotypic level (Fig. S9–10) showed this trend: drought-survival will increase only in the transition areas from Mediterranean and temperate regions (Fig. S9) and might decrease in areas that are already dry (Fig. S11). On the other hand, flowering time was predicted to decrease in the Mediterranean (Fig. S10–11). We note that the SNP effects on drought survival and flowering time were positively correlated, as disclosed by Canonical Correlation Analyses (section 3.4).

###### 4.2.4.4 Population genetics simulations

The predicted an allele in 2070 does not directly inform about the actual possibility of adaptation. This depends on (i) the frequency of the alleles and haplotypes in the population, (ii) the recombination rate, and (iii) the strength and efficiency of selection. Indeed, geographic predictions of alleles are probably bad predictors of allele frequency because random forest models tend to predict either one allele or another in a certain region. That is why we do not compare empirical present allele frequencies with frequency calculated from future predicted presence of alleles in different locations of Tübingen, Madrid and Malmö.

To obtain further insights into population dynamics required for adaptation, we simulated allele frequency changes in a Wright-Fisher population under a mutation-selection balance with inbreeding, as *Arabidopsis thaliana* is a selfer (code available at http://github.com/MoisesExpositoAlonso/popgensim). We started the simulations with the present frequencies of drought-related alleles of the 221 aGWA/GWA SNPs, with (codominant) selection coefficients (s) ranging from 0.01 to 20% fitness advantage. We considered SNPs as independent, that is, we did not include linkage disequilibrium information nor a recombination rate (see next section).

We carried out forward-in-time simulations for 50 generations, the approximate number of generations that natural populations of *A. thaliana* from today until 2070 at an average generation time of around 1.5 years (46). We assumed a mutation rate (*μ*) calculated from laboratory mutation accumulation lines (43, 47) and a selfing coefficient (*ψ*) of 99%, a conservative lower bound estimate from natural populations’ heterozygosity (48). The population size (*N*) was estimated from the genomic diversity in our dataset: The 40 genomes within Tübingen area had a genome-wide nucleotide diversity of 0.004 (section 3.1). With the equation:

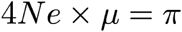
we solved for effective population size (*N_e_*) and transformed it into population size following the the relationship (49):

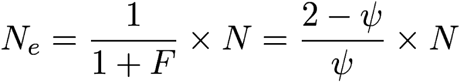

This yielded a N = 300,000 plants, which might be reasonable given that we consider an area of 50 km around Tübigen.

After running the simulations, we asked what selection coefficient would be needed to reach quasi-fixation frequencies of each allele (>0.9 frequency) or to match the drought-allele frequencies in Madrid or Malmö (assuming that those populations are adapted in comparison with the Tübingen one). When a specific allele frequency was higher in the Tübingen population than in Madrid or Malmö, we assumed selection would not be necessary and the coefficient was assumed zero. The results indicated that selection coefficients should be strong (but see (50)) for alleles to become fixed (Fig. S17). However, the distribution of selection coefficients were centered around 1-3% fitness advantage for Tübingen allele frequencies to match Malmö or Madrid (Fig. S17) (but see next section 4.2.4.5). We did not simulate different degrees of drift in our analyses as when the inequality: *N_e_s* > 1, holds, the weight of drift relative to selection is thought to be imperceptible (7).

###### 4.2.4.5 Considerations regarding recombination

As stated above, assessing whether a population can adapt depends on the frequency of drought-resistant individuals in the population, the rate of recombination to shuffle advantageous alleles, and the strength of selection. In our simulations above we did not work with haplotypes of SNPs in linkage disequilibrium as they are found in individuals, but considered SNPs to be independent. This can be seen *a priori* as a strong assumption. Simulations including whole haplotypes could inform about processes such as Hill-Robertson effect, hitchhiking, or background selection, which could be achieved with more complex approaches in the future (51).

Of relevance is that even in Tübingen there are already some individuals that already have many of the 151 GWA drought-resistant alleles, with one exceptional individual having 123/151 drought-survival alleles. The three next best individuals have 107, 105, and 99 alleles. Fifteen of the 28 drought-resistance alleles are not present in the Tübingen population and will have to be imported by migration. Therefore, to produce a hypothetical “fully adapted haplotype” with 136 alleles from the current standing variation (123 alleles are 90% of all 136 present alleles), only 13 drought-resistant alleles would have to be recombined and introgressed in the already advantageous haplotype. Such introgression events might not be limited by low frequencies of the advantageous alleles, as some were found at intermediate or as high as 90-95% frequency. Furthermore, in a scenario with a haplotype in the population with 123 alleles already present, simple individual-based simulations show that already with selection coefficients on the order of 0.5% advantage per allele, the 123 alleles haplotype will become completely fixed in the population within 50 generations (simulations not shown). Results of aGWA indicate similar patterns (24 alleles are 63% of all 32 present alleles) but more alleles are missing in the Tübingen population, as their frequency is lower and geographic distributions are narrower than GWA alleles.

We also used a series of approximate calculations to ascertain how many recombination events are required to generate a hypothetical “fully adapted haplotype”. In *A. thaliana*, there are on average 1.4 meiotic crossovers for each of the five chromosomes (52). Together with independent segregation of the five chromosomes, the parental haplotypes are rearranged at around 12 positions. A population of ~300,000 individuals (N) with a lower bound outcrossing rate of 1% (=1-F) over 50 generations (g), could thus undergo around ~2 million recombination events (= N * (1-F) * r * g), or about 1 event per 50 bp. Note that this could be a conservative estimate as rates of outcrossing in geographically close plants can be above 10% (53). This result suggests that recombination might be less limiting than expected *a priori.*

###### 4.2.4.6 GEM limitations

There is a long list of factors that we did not take into account and that will influence future plant response to climate change. We briefly enumerate them here:

A. We only focus on adaptation to drought, but other environmental stresses could have similar detrimental effects such as extremely high temperatures or ecosystem destruction. In addition, fluctuation in selection gradients and seasonal environmental variation are other possible consequences of climate change (54,55).
B. We only can explain ca. 50% of the drought survival variance with 221 SNPs.
C. We evaluated drought survival in a controlled greenhouse experiment, but the extrapolation to natural conditions may be difficult. This would require field experiments assessing fitness *in situ* to confirm that the identified SNPs actually report a fitness advantage (56).
D. Because the high narrow sense heritability suggests a mostly-additive genetic architecture, we carry out predictions with allele counts. However, we acknowledge that there is variation of the magnitude of the SNP effects (Table S3-4), and epistatic effects might exist.
E. All of our predictions are based on existing diversity, but de novo mutations are likely to make a contribution as well, especially in species with high reproduction rate, short generation time, and large population sizes (43, 57, 58).
F. Biotic interactions can also play a relevant role of population dynamics, which we ignored (59–61).
G. In addition, although long-term evolution should be driven by genetic adaptation, it is expected that phenotypic plasticity will partially buffer the detrimental consequences of environmental change (62).
H. The existence of a seed bank in *A. thaliana (46, 63)* would cause longer generation times and overlapping generations, and both alter the speed and dynamics of allele frequencies (64).
I. Although our rough calculations suggest that recombination would not be a limitation for future adaptation in *A. thaliana* populations, we have not incorporated such processes in our modeling, as it is not a trivial matter (51, 65). This ignores phenomena such as background selection or hitchhiking effects that could arise from phenotypic trade-offs and the currently realized composition of haplotypes in the population.

## SUPPLEMENTARY FIGURES & MEDIA

**Figure S1.**
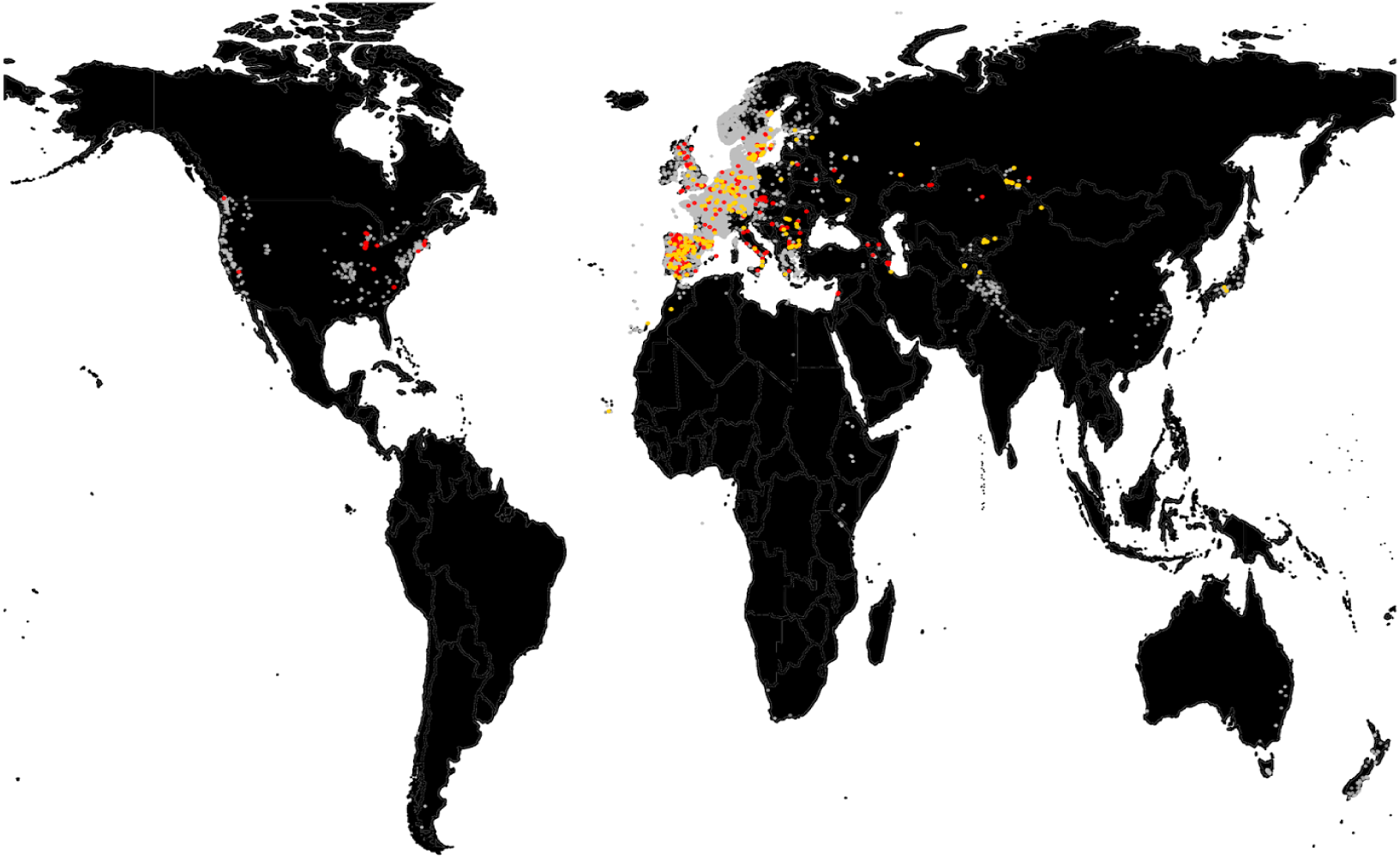
Extent of *Arabidopsis thaliana* distribution. The global map shows ca. 80,000 records from the Global Biodiversity Information Facility (GBIF, www.gbif.org) (grey), the 762 global accessions used for genetic analyses (red), and 210 accessions used for phenotyping experiments (yellow).

**Figure S2.**
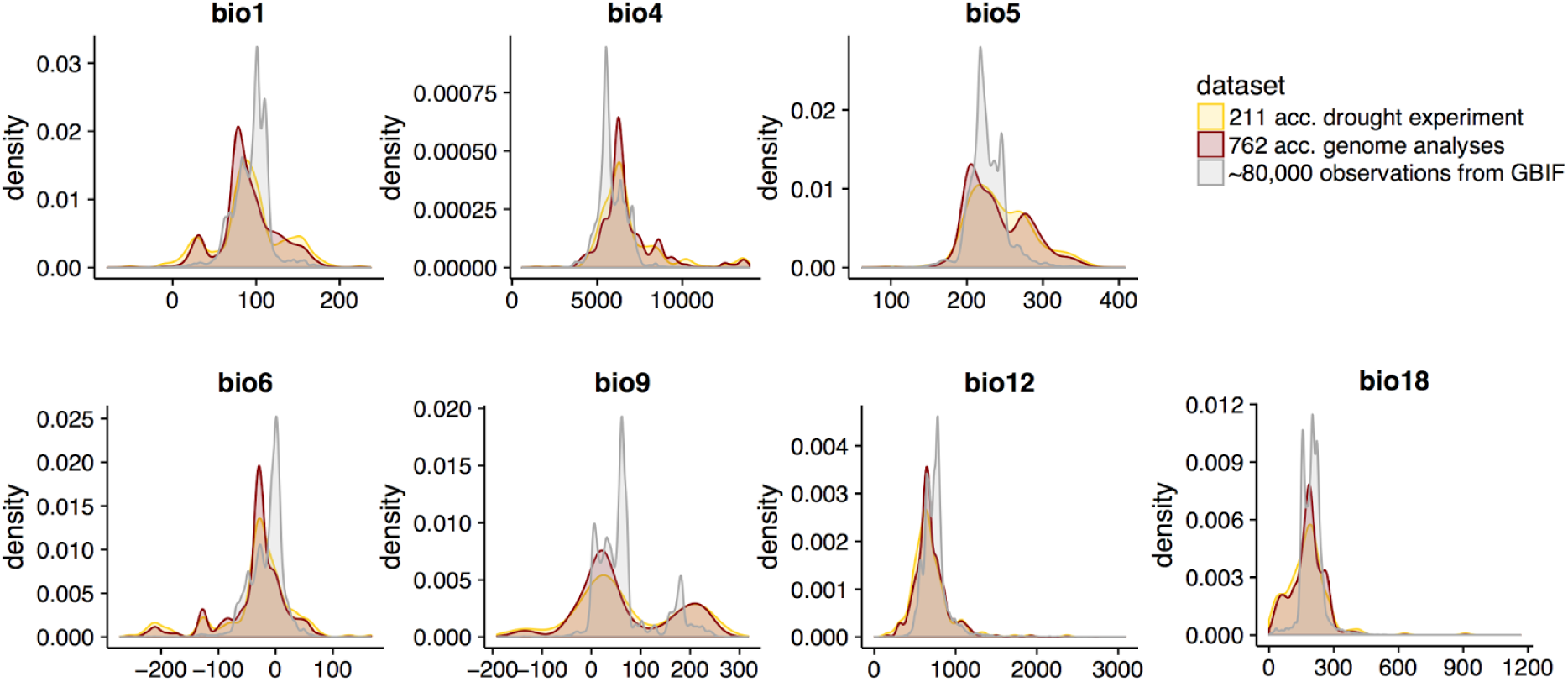
Environmental ranges of *Arabidopsis thaliana*. We show the range in key environmental variables for the three datasets in Fig. S1. The set of accessions used in our analyses not only covered the range of the species as estimated from GBIF data, but also showed that these accessions have a more even distribution throughout the environmental ranges. The bioclimatic variables are: annual precipitation (bio12), precipitation of the warmest quarter (bio18), annual mean temperature (bio1), temperature seasonality (bio4), maximum temperature of the warmest month (bio5), minimum temperature of the coldest month (bio6), and mean temperature of the driest quarter (bio9). See Table S5 for more information.

**Figure S3.**
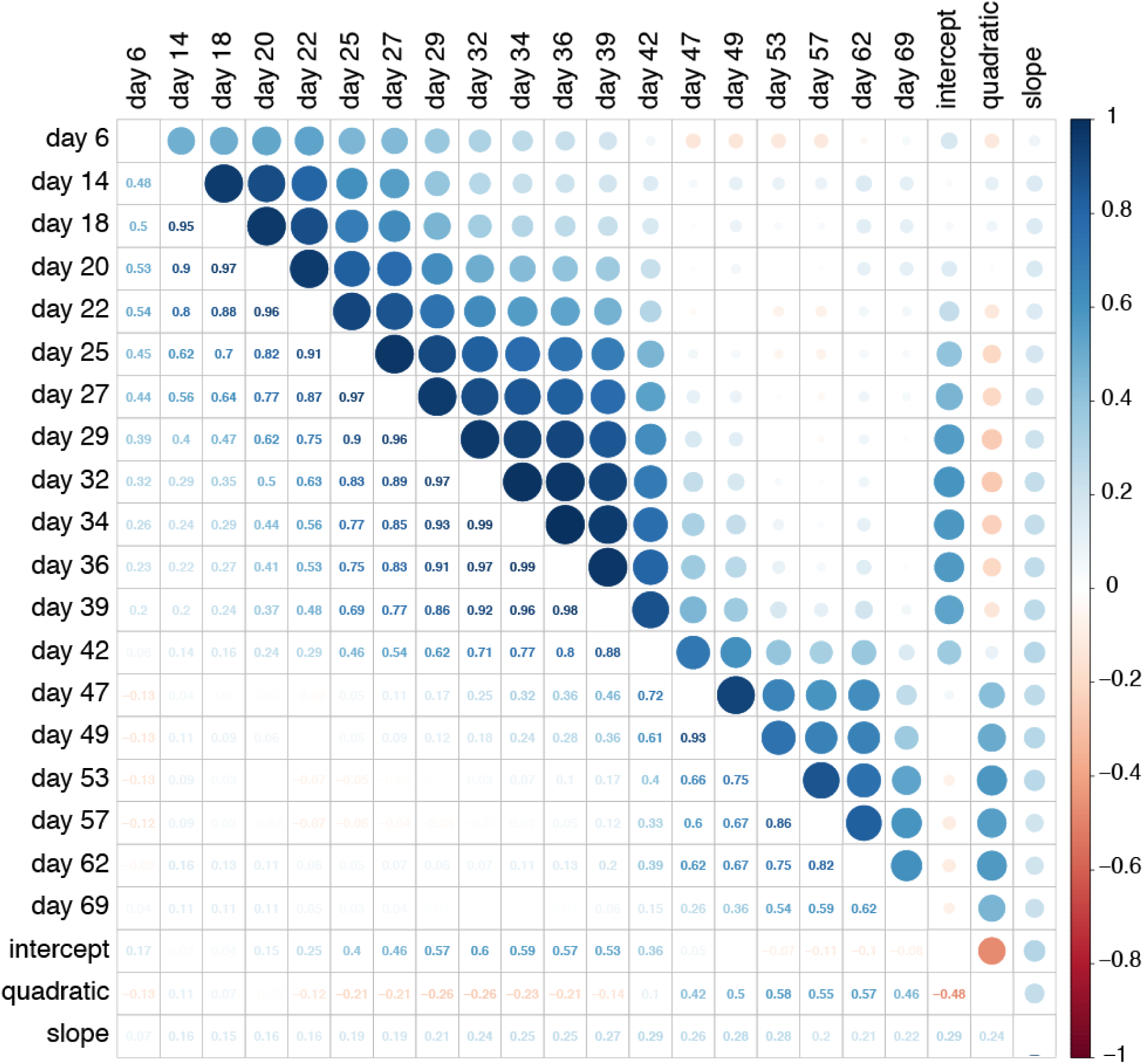
Correlation between raw green pixels in plant images and model parameters. Pearson product-moment correlation coefficients between the three drought-index parameters and the ‘raw’ number of green pixels per pot. Sizes of circles indicate strength and colors signs of association, shown as numbers in the lower triangle.

**Figure S4.**
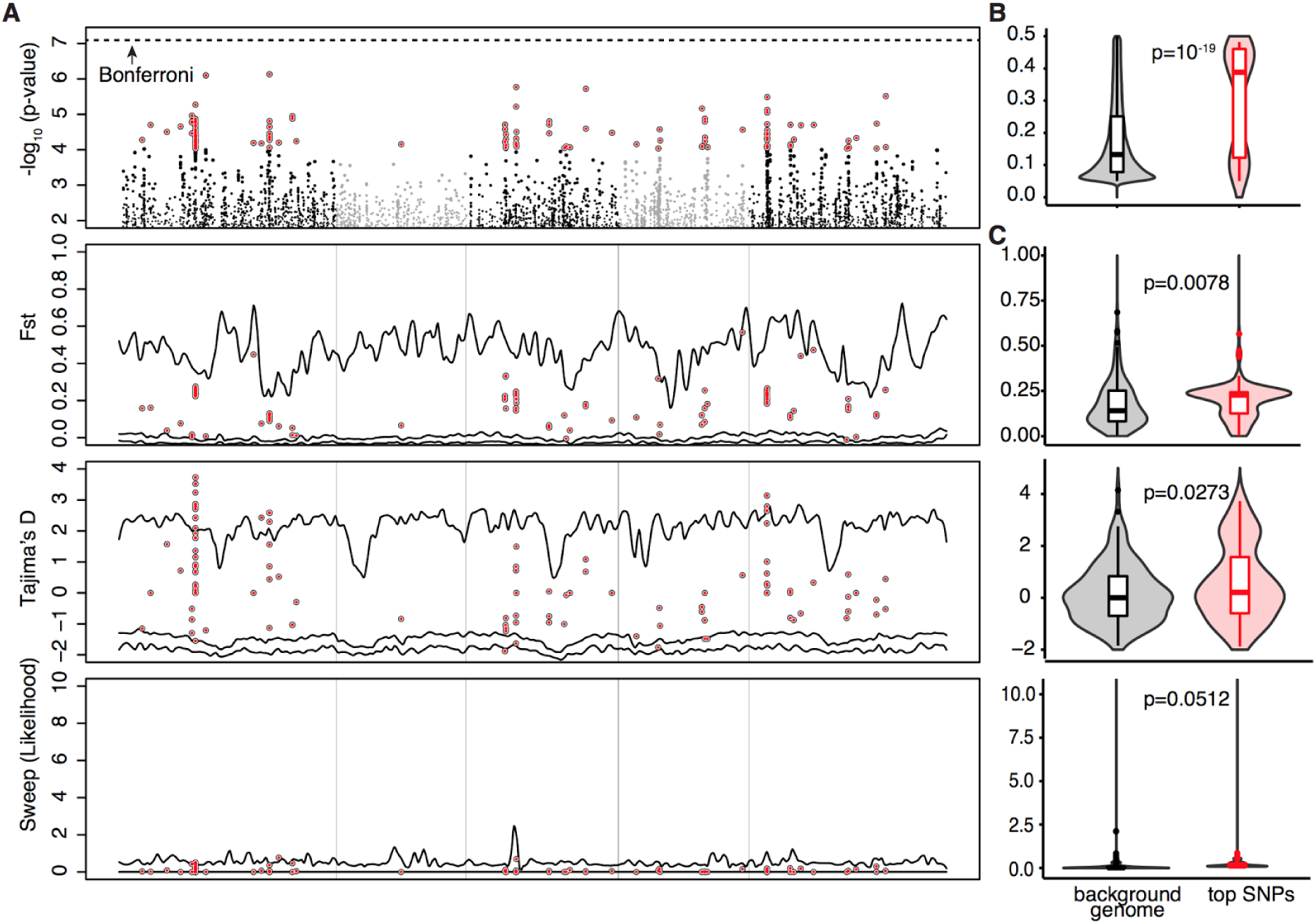
GWA with drought survival and population genetic statistics. (**A**) Manhattan plot of drought resistant GWA, F_st_, Tajima’s D, and selective sweeps. (**B**) Violin and box plots of allele frequency, and (**C**) F_st_, Tajima’s D, and selective sweeps of the top 150 SNPs (red) vs frequency-matched 150 SNPs from a random genome background (grey). GWA was calculated using EMMAX. F_st_ across populations (see Fig. 1) and Tajima’s D were calculated using PUNK. Sweep likelihood was calculated using SWEED software. Median p-values from Wilcoxon tests with 100 bootstrap replicates are indicated.

**Figure S5.**
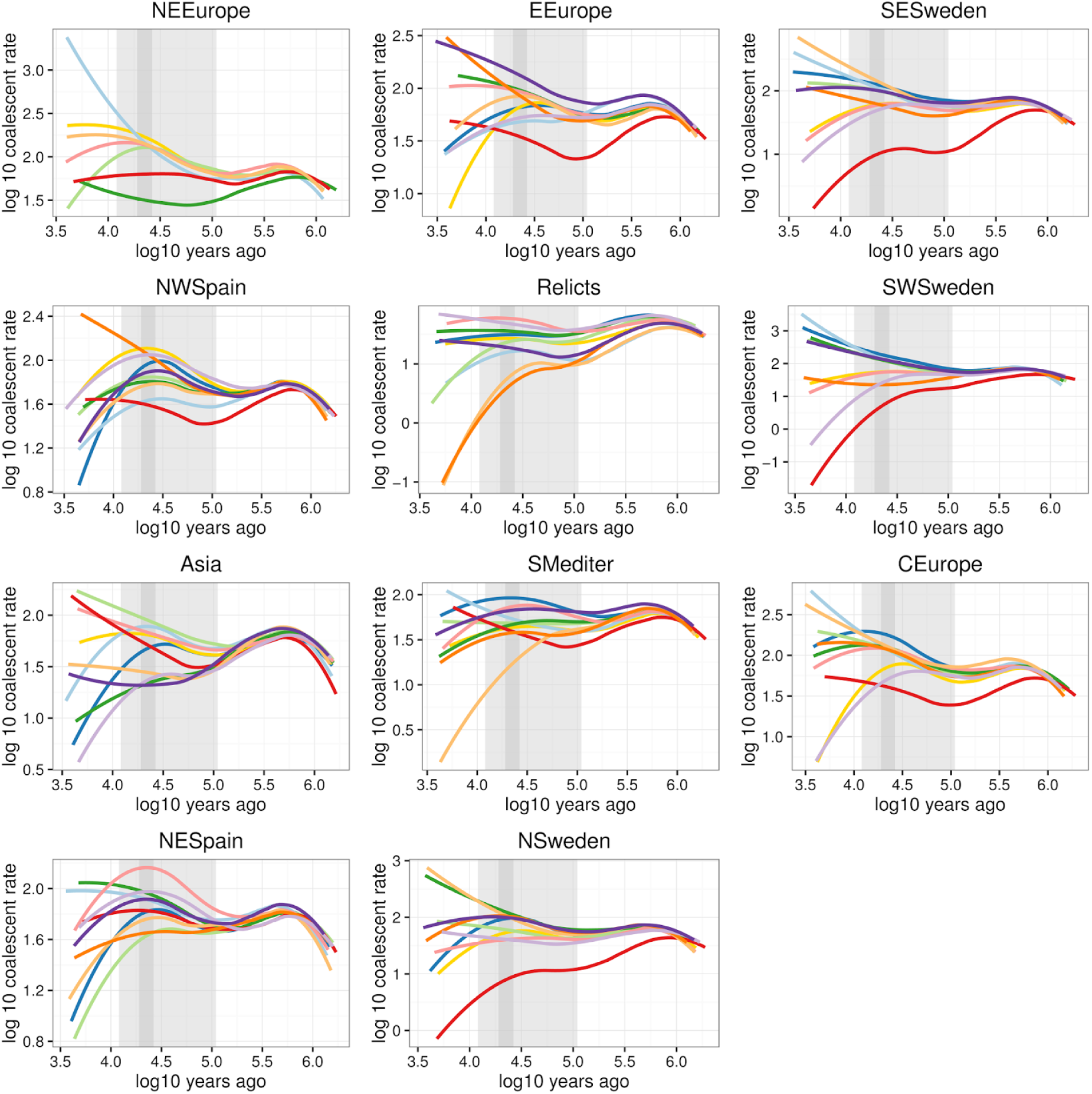
Cross-coalescent rates between populations inferred by MSMC. Joint coalescent rates of each of the 11 ADMIXTURE genetic groups are (see Fig. 1 and Fig. S12) compared to the other groups. Each line is a smoothed loess of 6 replicated runs. Light grey area indicates the extent of the last glacial maxima (100-10 kya) and dark grey area the peak of the last glaciation (22 kya). Analyses between certain groups failed (e.g. NE Europe with Asia), likely due to proximity between genomes. Note that the N. Swedish group is the first to separate from the relicts.

**Figure S6.**
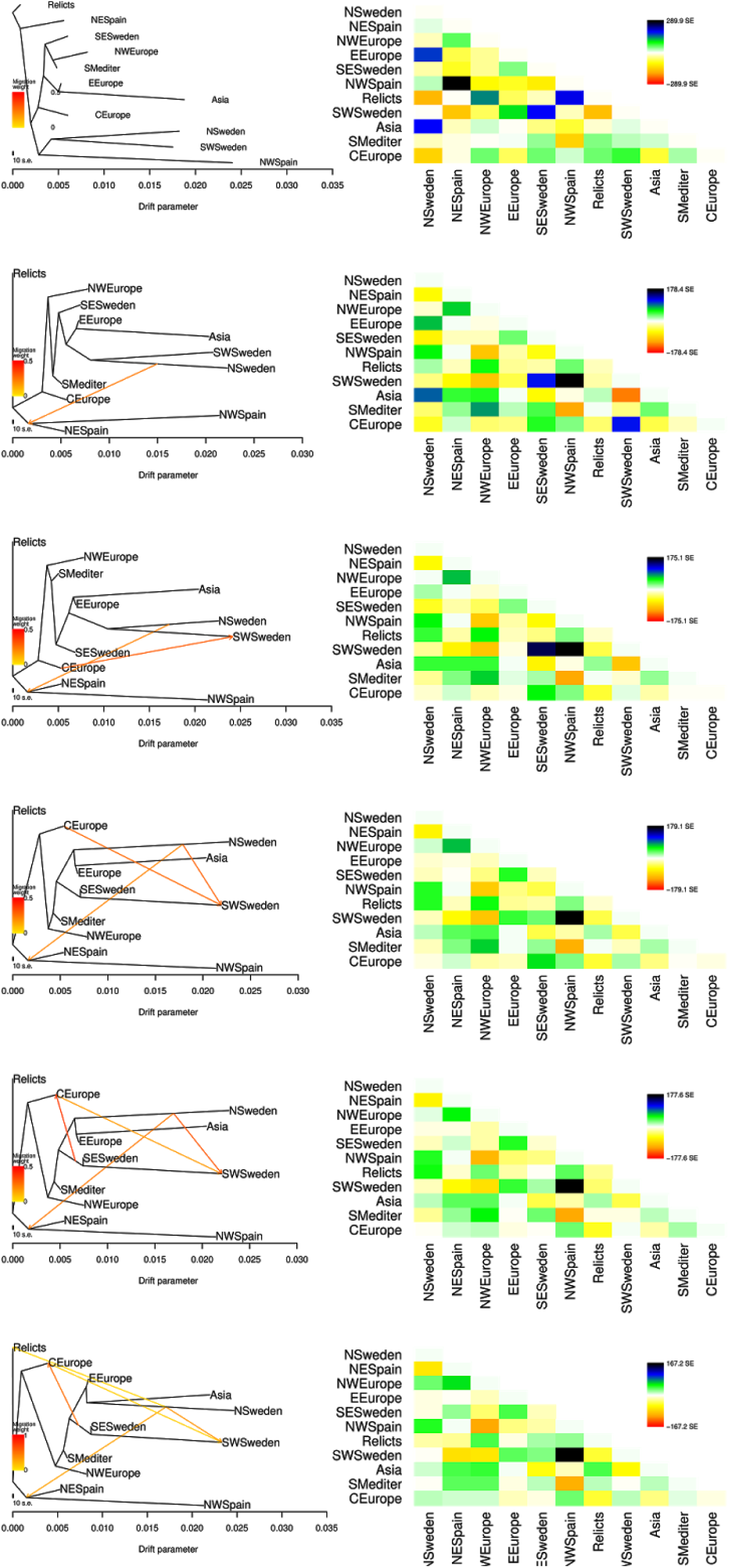
Treemix with different migration rates. Maximum likelihood (ML) population trees from Treemix (left). Analyses with zero to five migration edges are presented. Heatmaps with the residual fit of the ML trees are shown on the right. Note that the unexpected closeness of NW Spain and Sweden without migration is resolved with one migration edge. With this, a more parsimonious tree that adheres to geographic locations is uncovered.

**Figure S7.**
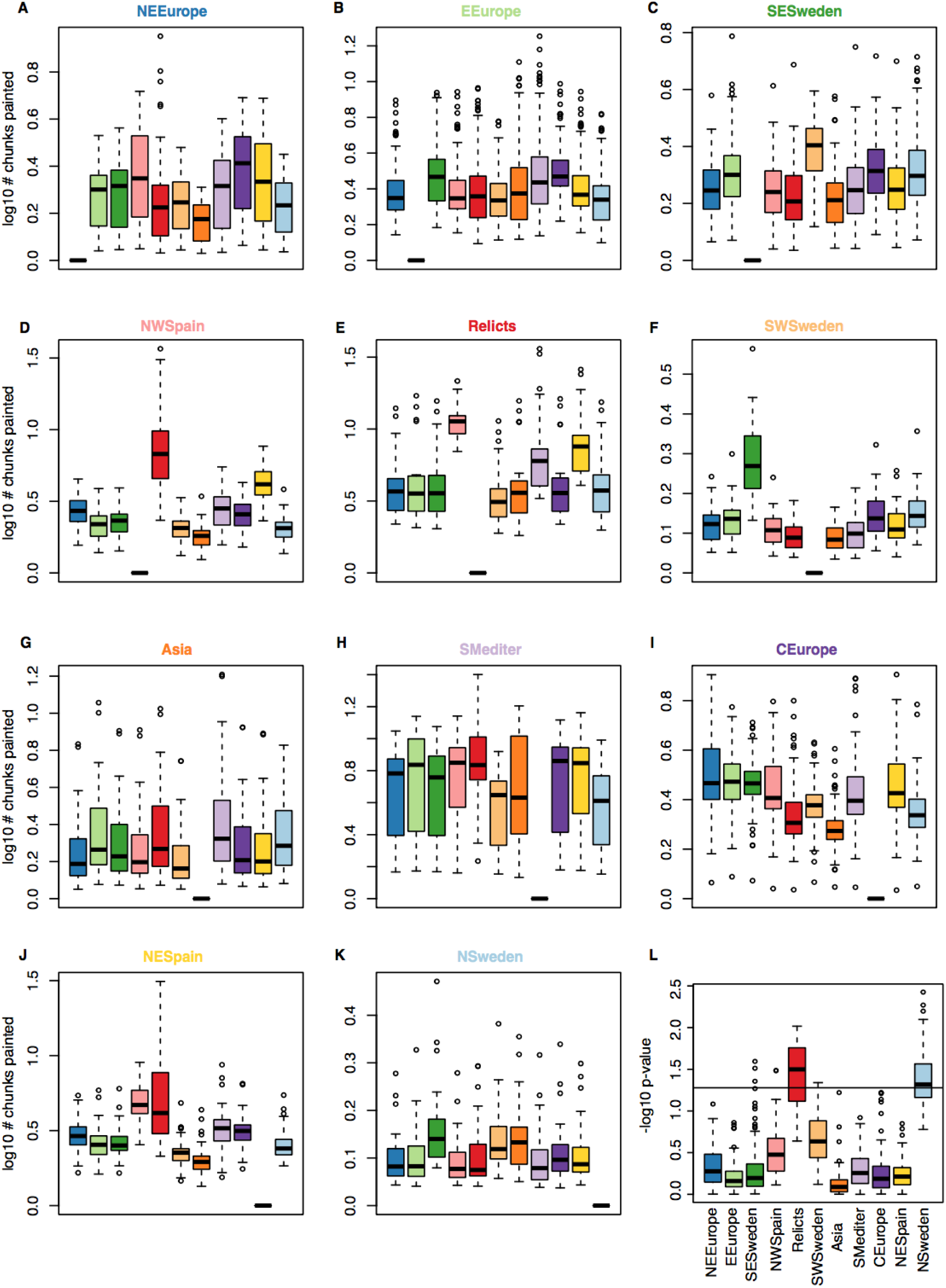
Genomic ChromoPainter chunks per population. **(A-K)** Summary of the number of ChromoPainter chunks inherited from other genomes that had been assigned to ADMIXTURE groups. Each graph summarises the information of all the genomes from an admixture group. (**L**) The p-value of the Pearson correlation test between an accession’s drought survival index and the number of chunks received from another genome. The p-value distributions of genomes from the same ADMIXTURE group are grouped in a box plot. Intuitively this can be interpreted as how well the number of chunks inherited from a specific donor predicts the drought survival of the receiver. The black line indicates the 5% significance threshold, which is passed by most relict and N. Sweden groups. Therefore, chunks that have N. Swedish and/or relict ancestry explain the drought survival of other individuals well.

**Figure S8.**
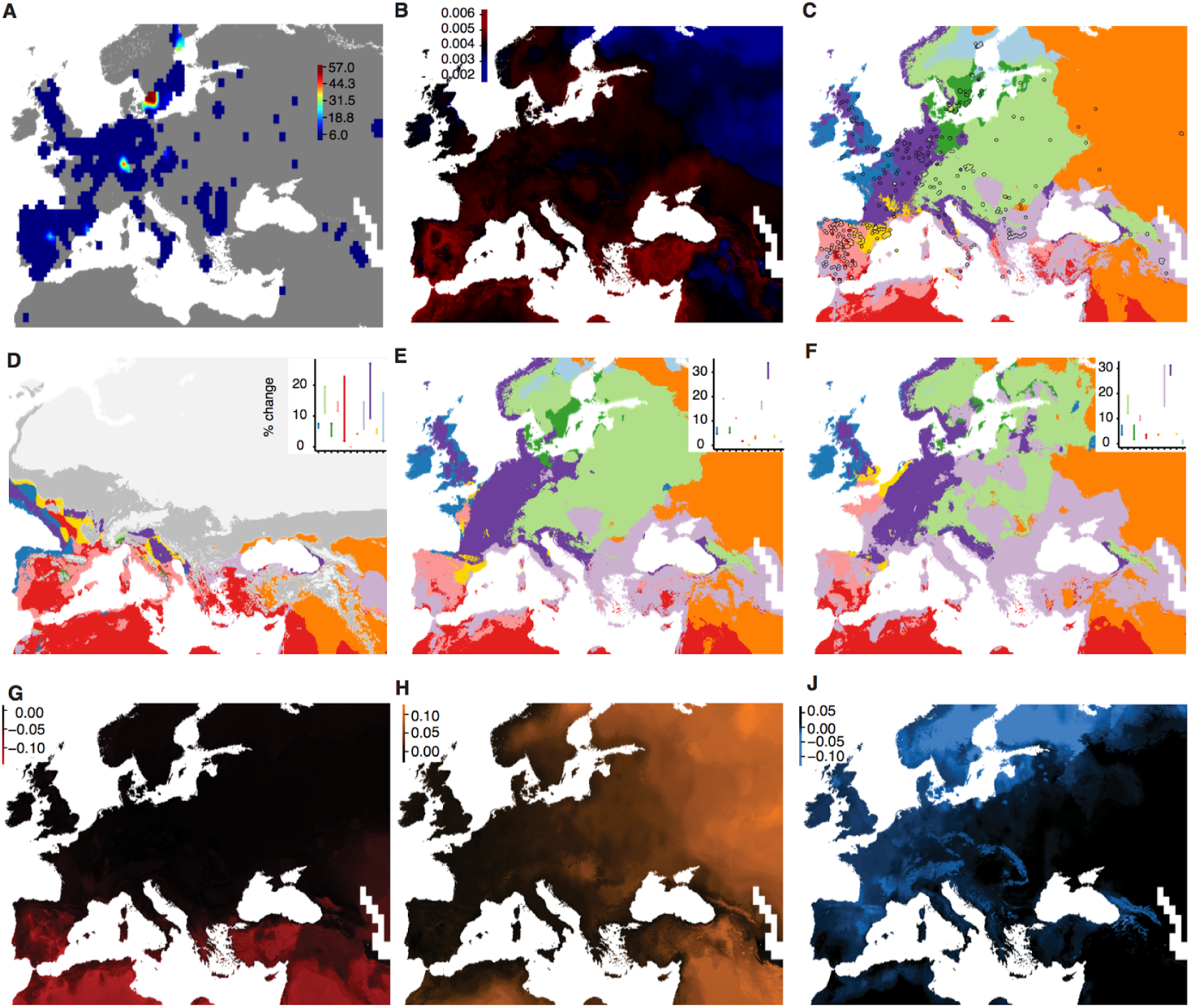
Environmental niche model of genetic diversity and population structure. **(A)** Distribution of 762 accessions from the 1001 Genomes project used for environmental niche modeling of genetic diversity and analysis of population structure. Colors indicate number of accessions within a 1°×1° latitude × longitude grid. **(B)** Random forest environment niche models using estimates of pairwise nucleotide diversity (π) diversity of each accession with its closest 10 geographic neighbours. The trained model was used to predict diversity based on environmental data. **(C)** Random forest environment model of the 11 genetic groups (see Fig. 1). Locations with accessions are shown as points filled with the actual genetic group assigned, and are used for model training as in (B). The trained model was used to predict a raster of environmental variables and is shown in the background. When the circle is filled with the same color as the background, the model succeeds in the prediction. The trained model was also used to predict presence of different genetic groups at the Last Glacial Maxima **(D)** and for 2070 under low **(E)** and high **(F)** CO_2_ concentration scenarios, (see Fig. 1 for color keys). **(G-J)** The first three genome-wide principal components from Fig. 2 were modeled based on environmental variables. Later, these were used as covariates of Genome-Environment Models (Fig. S13–14).

**Figure S9.**
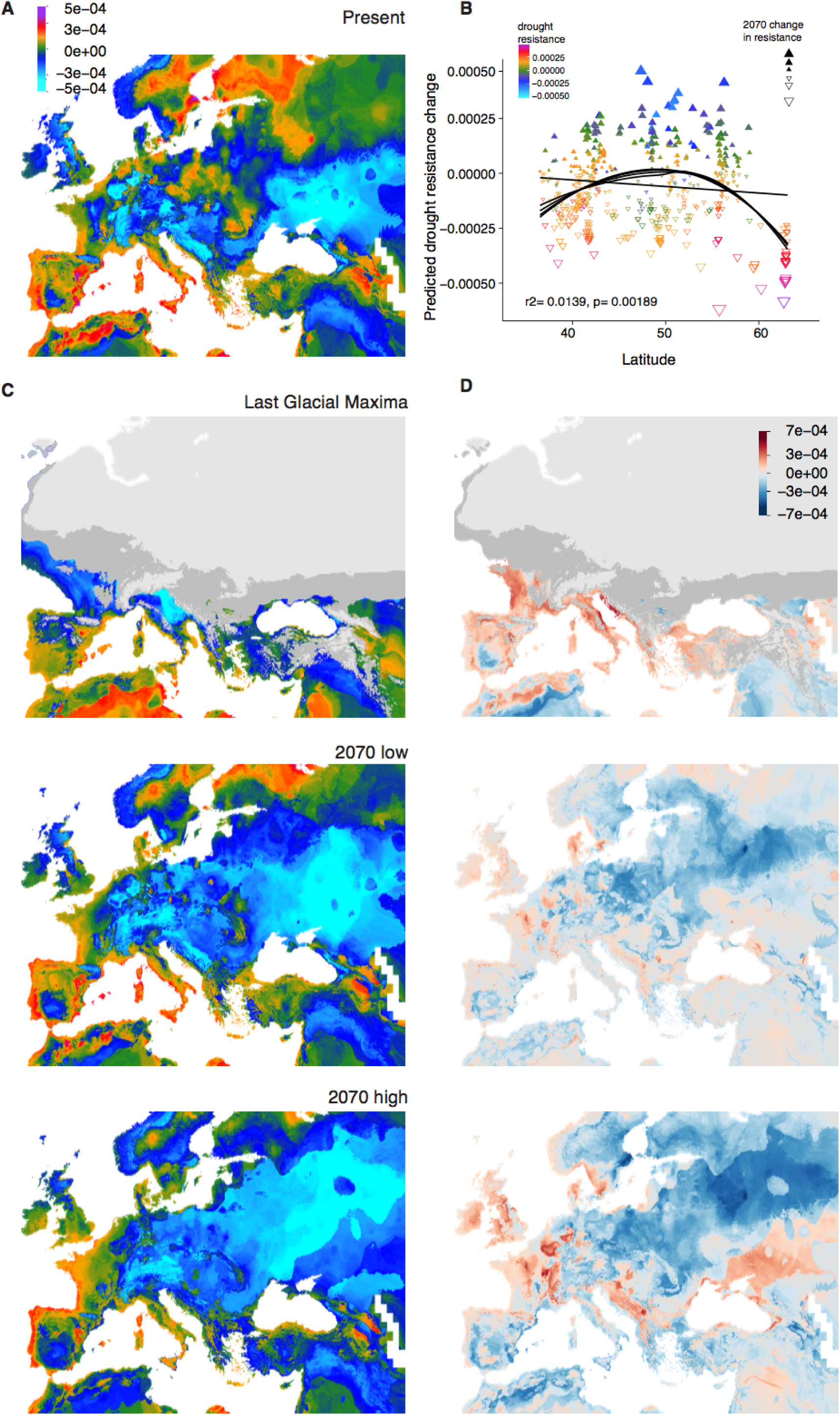
Environmental niche model of drought index. **(A)** Present geographic prediction of drought survival index from a random forest environment niche model trained on experimentally determined phenotypes for 211 accessions. Note that the highest drought survival values are inferred for the Mediterranean as well as N. Sweden. **(B)** Correlation of phenotypic change in 2070 under a high CO_2_ scenario with latitude; colors indicate present drought survival values. **(C)** The trained model is also used to predict drought survival index under the Last Glacial Maximum, and for two 2070 scenarios of low and high CO_2_ concentrations. **(D)** For the three scenarios, the change is shown relative to the current date prediction for easier comparison.

**Figure S10.**
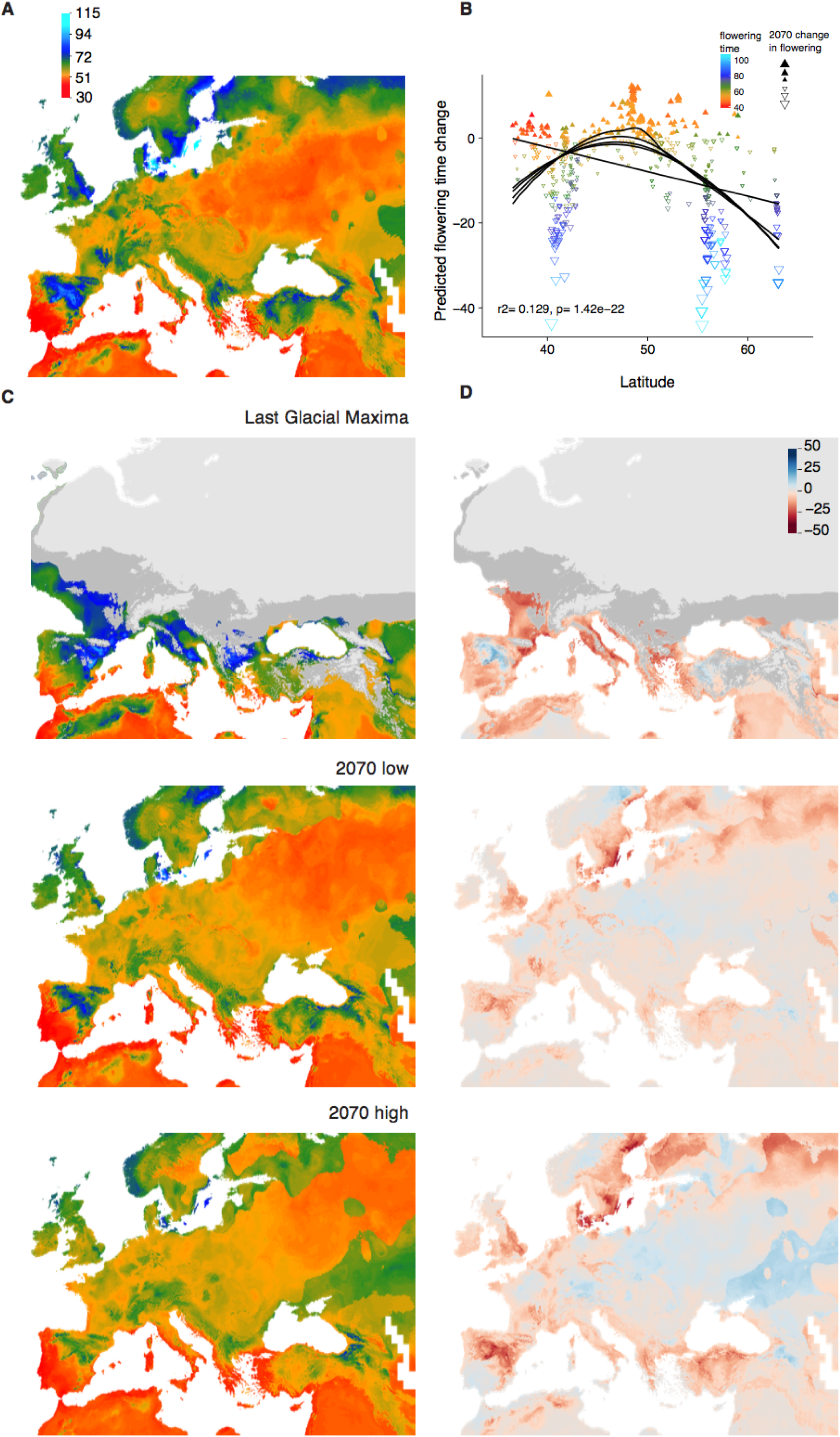
Environmental niche model of flowering time. Same models as in Fig. S9, but for flowering time.

**Figure S11.**
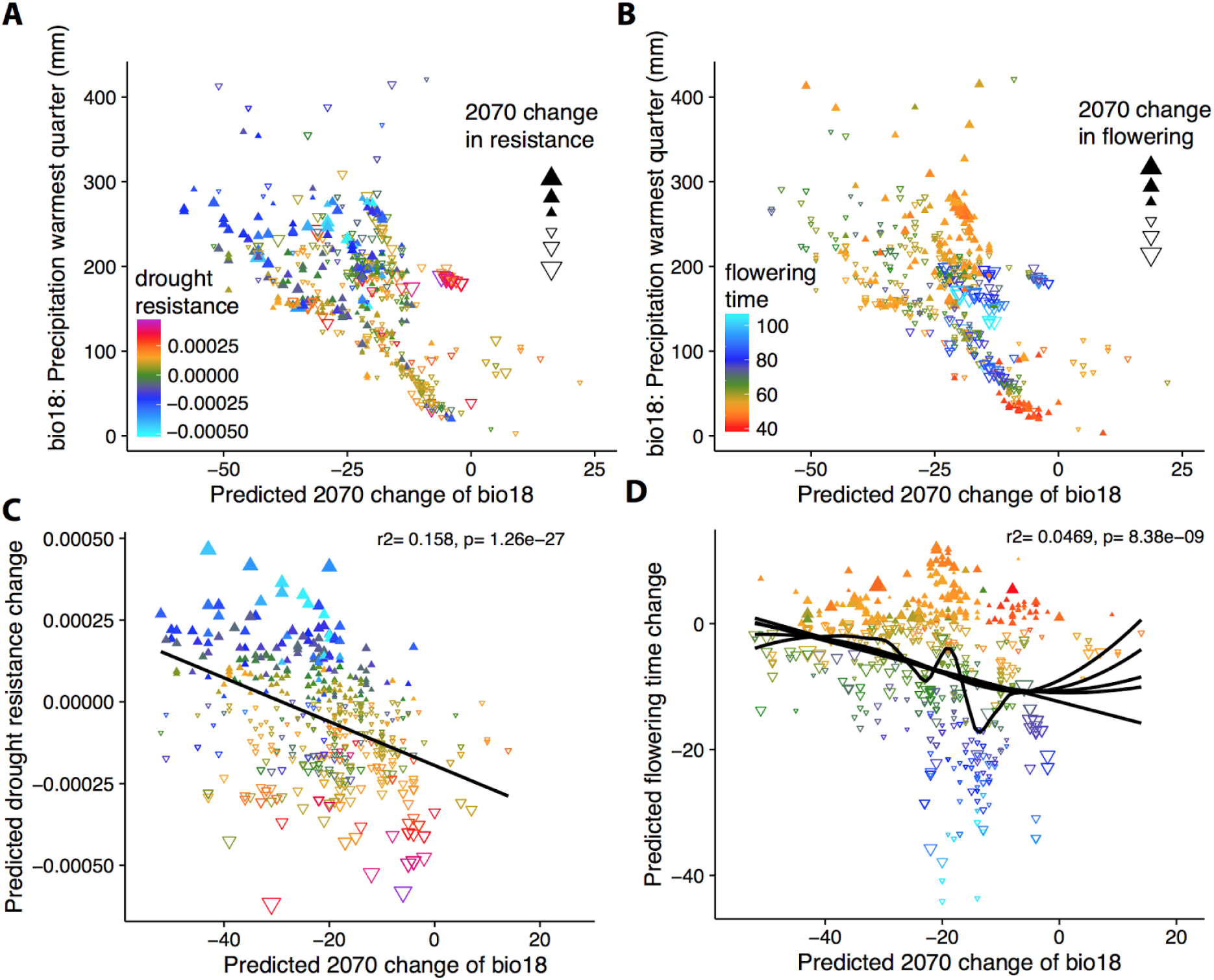
Profile of phenotypic change under climate change. **(A, B)** Correlation of precipitation during the warmest quarter today and in 2070 under a high CO_2_ scenario. Colors indicate current drought survival **(A)** or flowering time **(B)**, and shapes indicate increase or decrease in trait values for 2070. **(C, D)** Regression of the predicted change in drought-survival **(C)** and flowering time **(D)** on the predicted change in precipitation in 2070. Note that areas with already low precipitation will not have large decreases in precipitation in 2070 (A-B). Note also the linear relationship between decreased precipitation in 2070 and predicted increase of drought-survival in (C). Flowering will be on average faster in 2070 (D), but the relationship between precipitation reduction and flowering time change is not linear, which suggests that areas with a moderate reduction in precipitation will have accelerated flowering (rather than increased drought survival).

**Figure S12.**
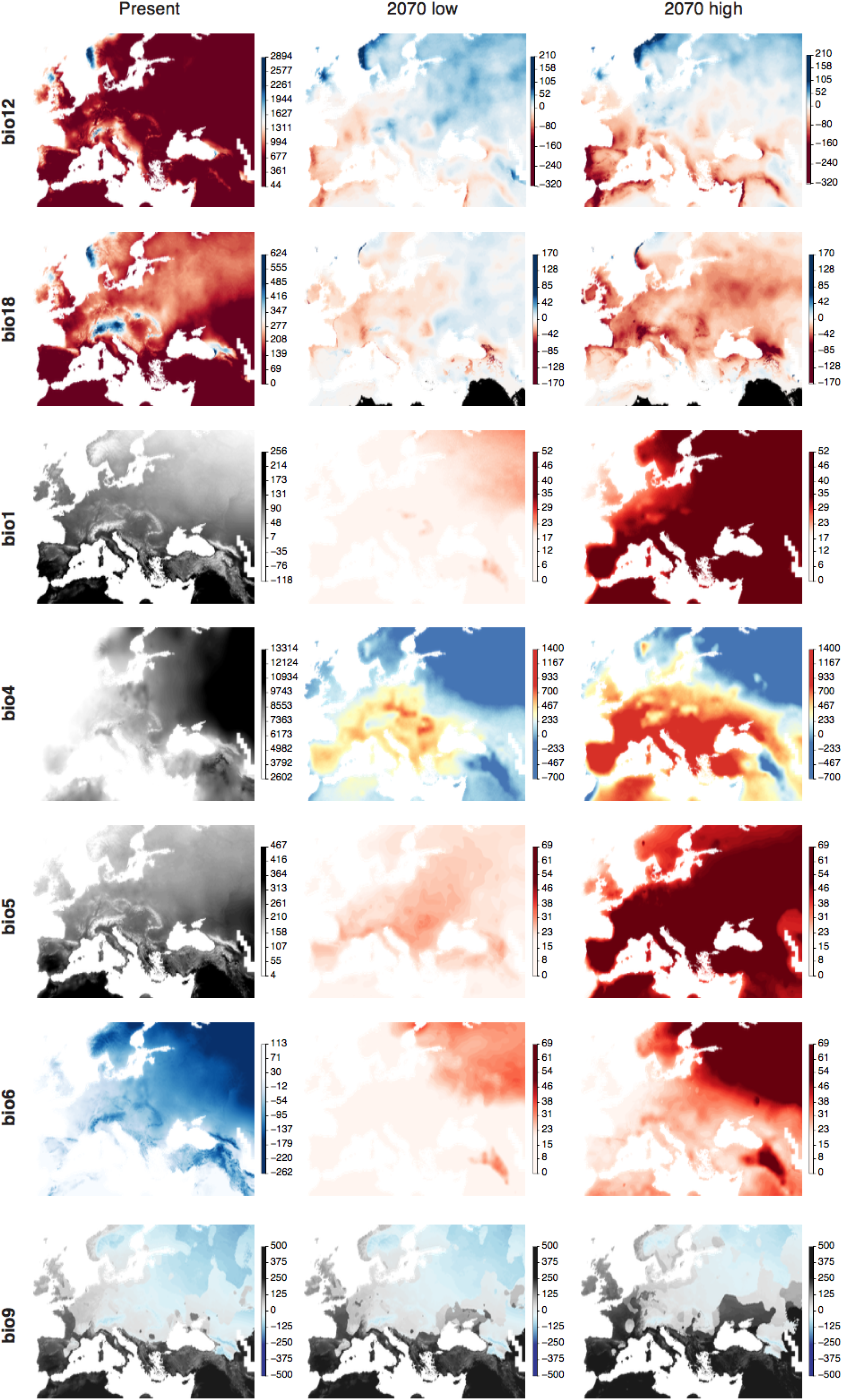
Maps of the most important climatic variables. The bioclimatic variables (www.worldclim.org) that typically had more importance in phenotypic and genome environmental models are shown as an aid for interpretation of the results from our study, bioclim variable shown are annual precipitation (bio12), precipitation of the warmest quarter (bio18), annual mean temperature (bio1), temperature seasonality (bio4), maximum temperature of the warmest month (bio5), minimum temperature of the coldest month (bio6), and mean temperature of the driest quarter (bio9). The columns show distributions at present, in 2070 under a scenario of low CO_2_ concentration, and in 2070 under a scenario of high CO_2_ scenario. Except for bio9, the values for future scenarios were expressed as future-present difference to highlight geographic areas that will change the most. Note the bimodality of bio9: areas in black are summer drought (Mediterranean climate) areas, whereas blue areas are winter-drought. Also note that bio18 is predicted to change mostly along the transition from the Mediterranean to non-Mediterranean climate. In bio18, areas that will be under lower precipitation than any current location of *A. thaliana* are shown in black, to highlight that most areas will remain within the range of current precipitation across the species range.

**Figure S13.**
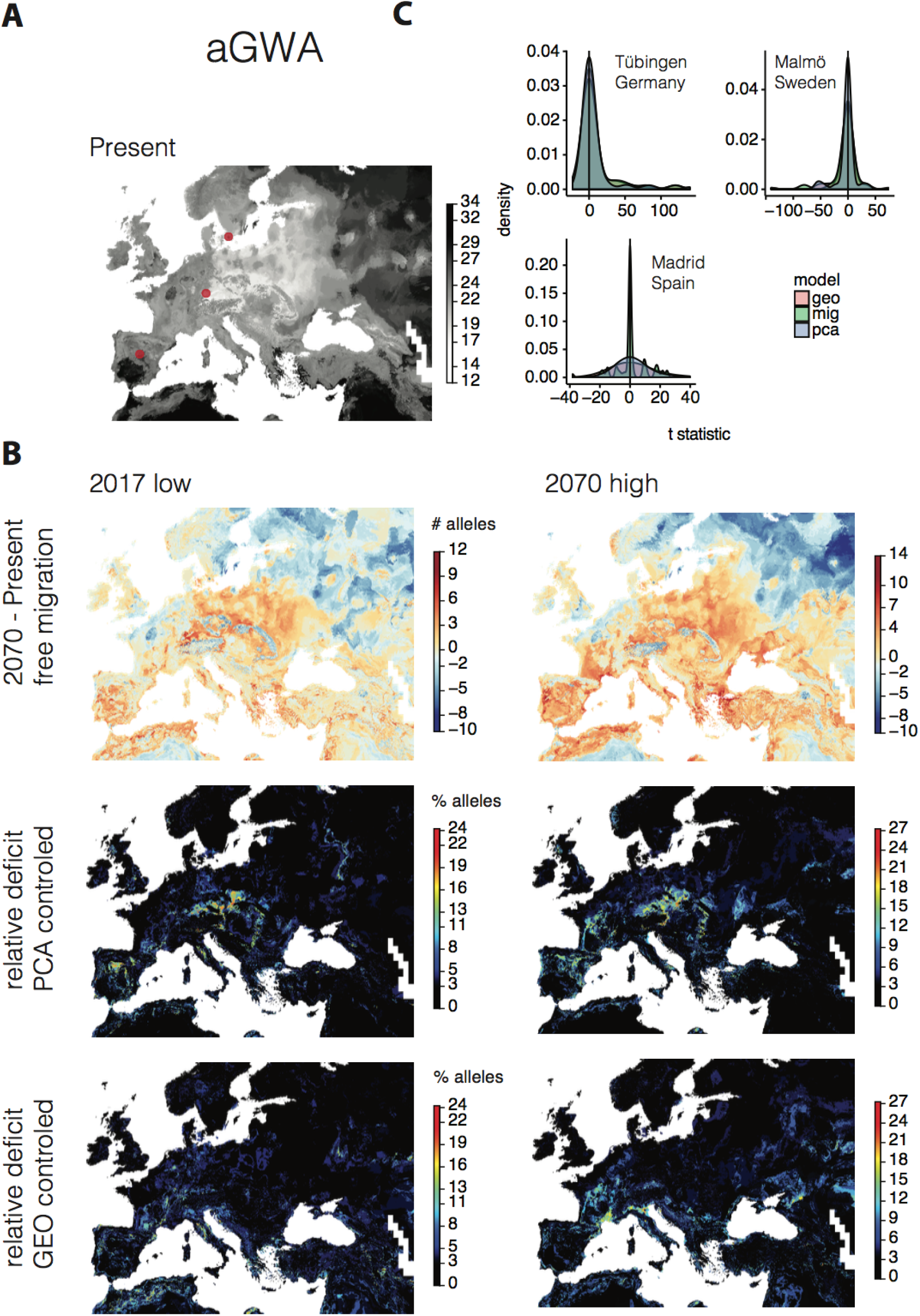
aGWA Genome Environment Models (GEM) **(A)** We ran GEMs to describe the geographic distribution of alleles at the 70 aGWA top loci. Concatenating all maps, we produced a map of the count of all drought-survival alleles that a genotype is expected to have in a given location today. **(B)** The trained model from (A) was used to predict distribution of drought alleles in the future. The difference to numbers inferred for today (A) corresponds to the alleles that will have been gained or lost in 2070 in a given location. Two additional models were trained which included a genome background (PCA) correction and latitudinal and longitudinal (GEO) correction of the allele distributions. The percentage of gained alleles from the “free” model that were not present in the corrected models is shown as a deficit in percentage. **(C)** For three highly sampled locations, Madrid (Spain), Tübingen (Germany) and Malmö (Sweden), we calculated allele frequency differences between today and 2070 (under high CO_2_) and calculated a t-statistic to describe the effect size of the change. A skew towards the right (increase) is observed for Tübingen only.

**Figure S14.**
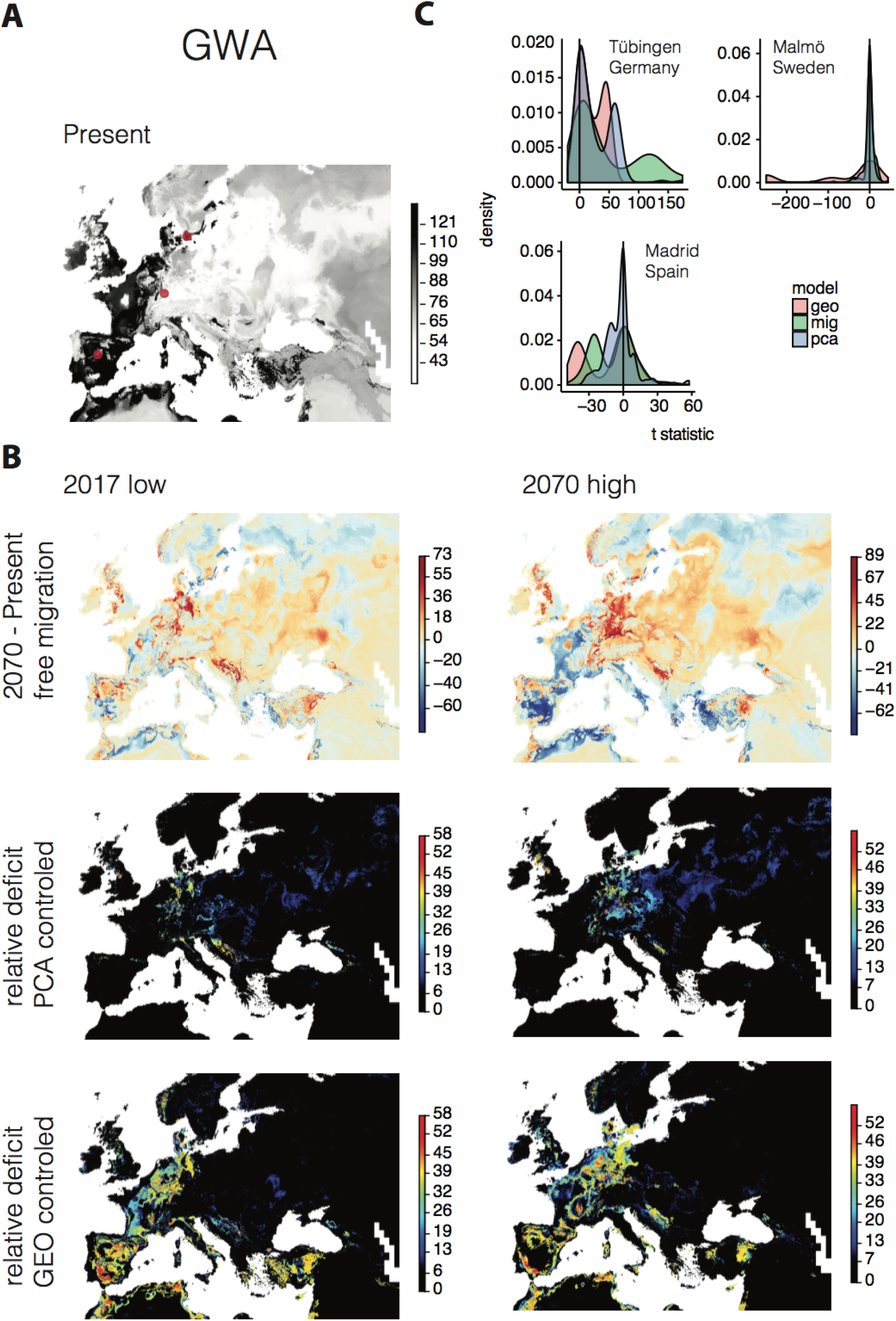
GWA Genome Environment Models (GEM) See Fig. S13 for legend.

**Figure S15.**
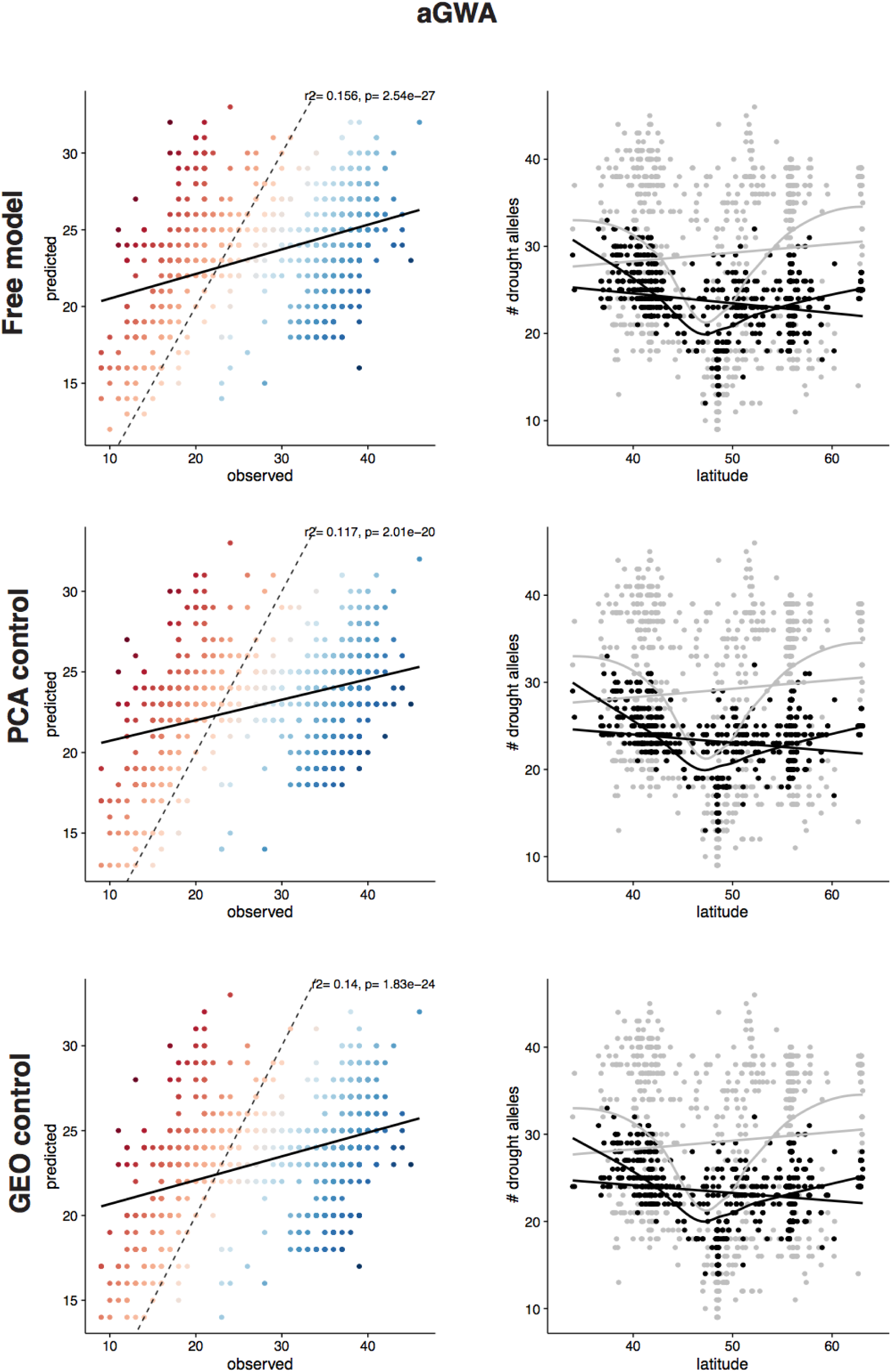
aGWA GEM residuals. For each GEM, we plot the predicted against the observed (empirical) number of drought-associated alleles at each sampled locations. Red color indicates overestimation and blue underestimation. Latitudinal trends of predicted (grey) and observed (black) are shown (right). Note that the variance of predictions is larger than the empirical observations, probably due to the discrete nature of random forests.

**Figure S16.**
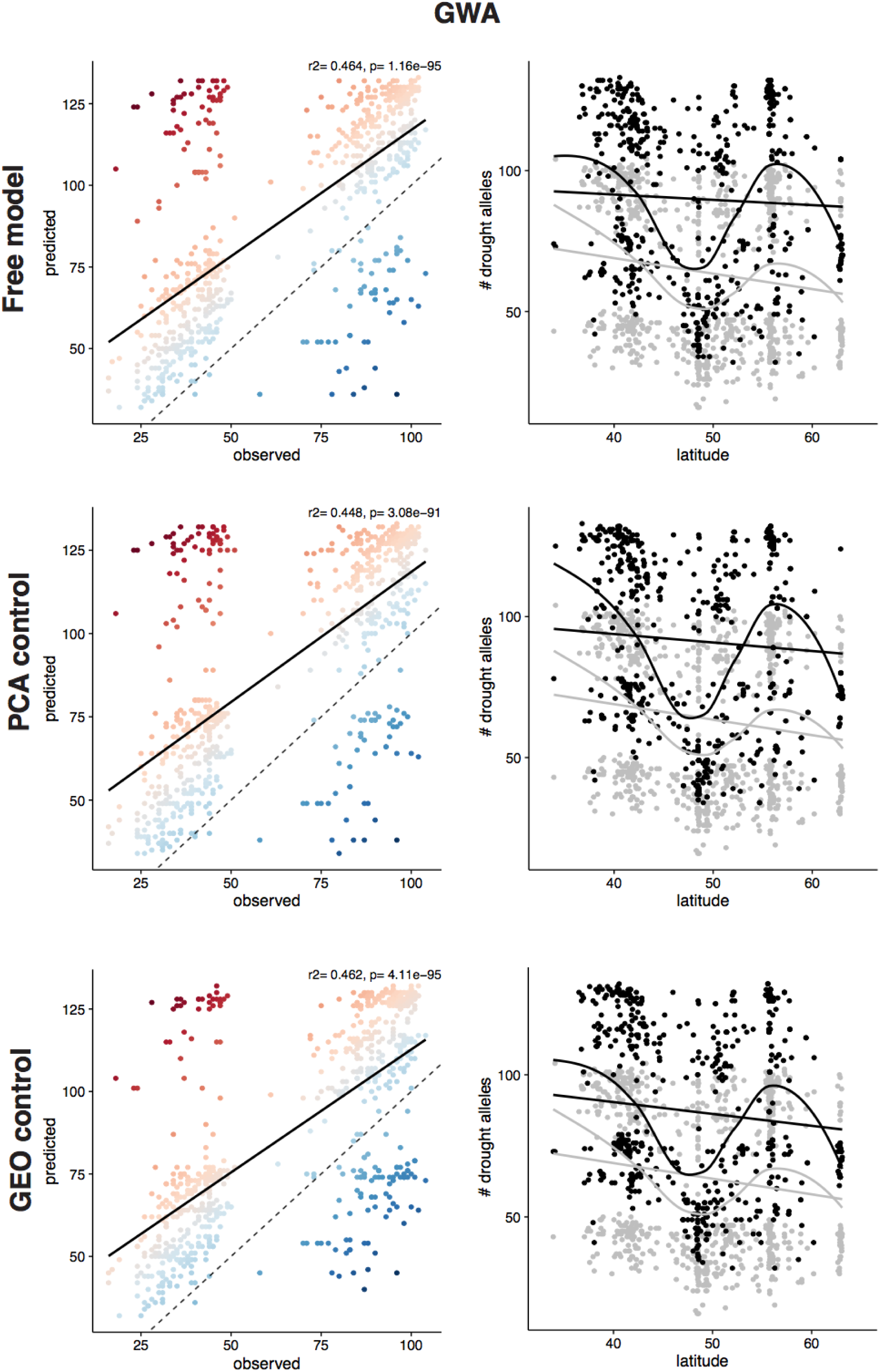
GWA GEM residuals. See Fig. S15 for legend.

**Figure S17.**
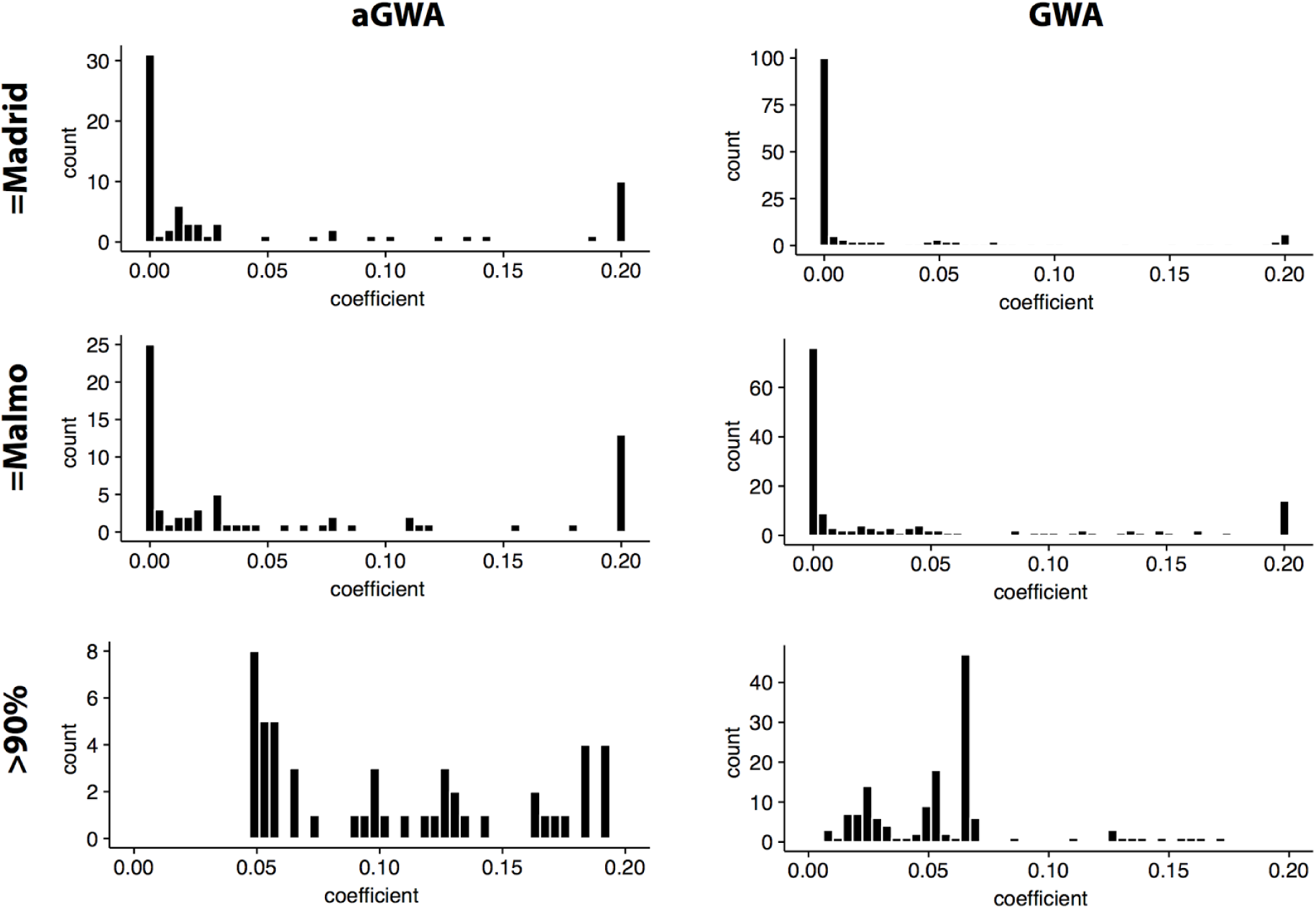
Population genetics simulations. We ran Wright-Fisher population simulations of 70 (aGWA) or 151 (GWA) independent loci for 50 generations of evolution under mutation-selection balance, starting with the current allele frequencies in the Tubingen population., and repeating each simulation with an array of selection coefficients from 0.0001 to 0.2 (relative fitness advantage) for each locus. The distributions shown correspond to the positive selection coefficients that are required for the drought-alleles to rise to the frequency at which they are currently found in Malmö (top) or Madrid (center), or to at least 90% (bottom), which is close to fixation.

**Video S1.**
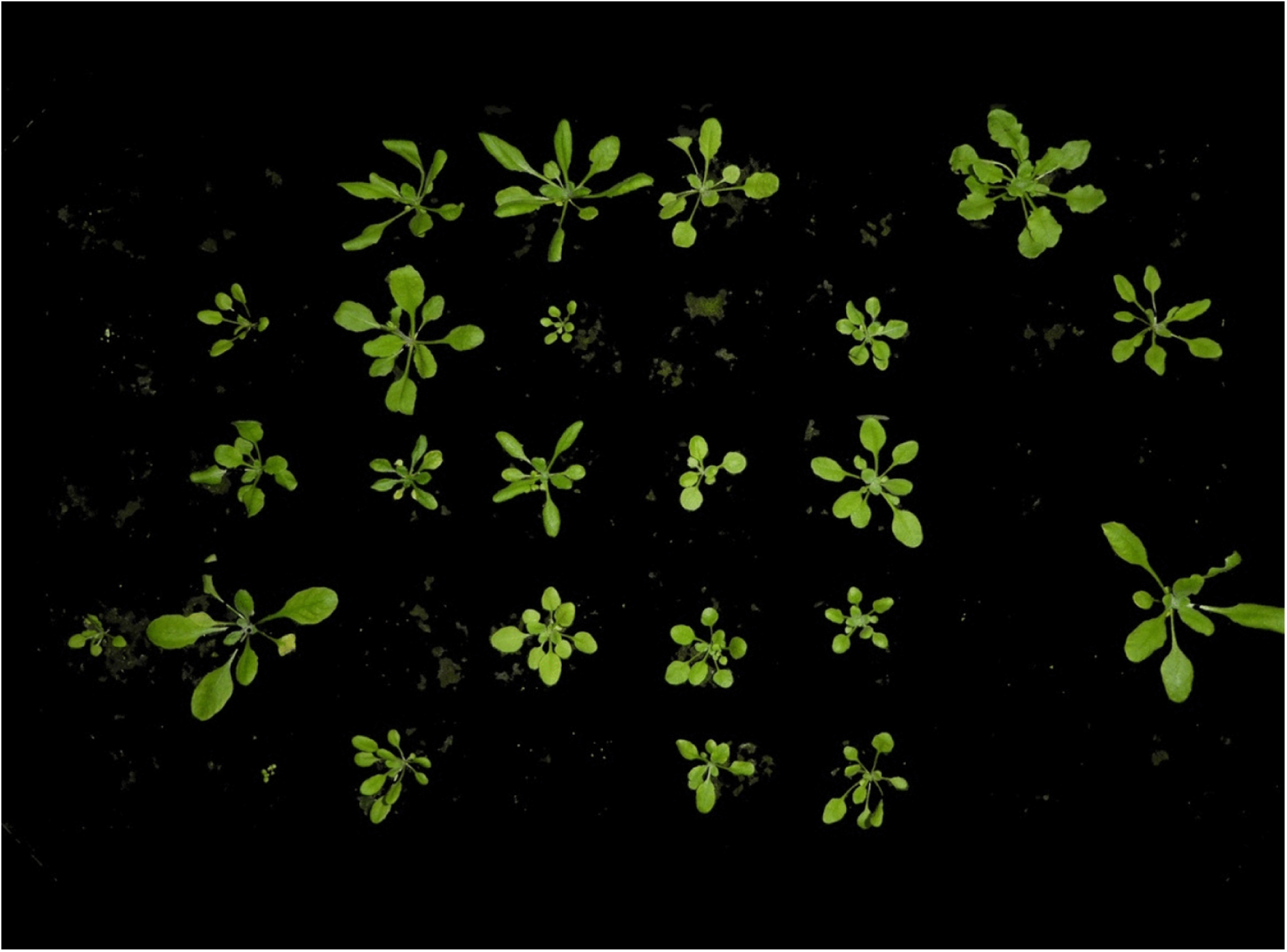
Example of segmentation. 19-frames time series of green-segmented images for one exemplary tray. See the online file Video_S1.gif (or click here).

## SUPPLEMENTARY TABLES & DATASETS

The combined tables can be downloaded as.xlsx file from: http://github.com/MoisesExpositoAlonso/Exposito-Alonso_2017_drought_Supplementary_Table_s [*here link to the journal’s web upon publication*]

**Table S1. Accession information.**

1001 Genomes IDs, common names, countries of origin, and geographical and environmental information.

**Table S2. ADMIXTURE cross-validation** for all possible groups from 2 to 20.

**Table S3. Selection signatures and annotation of top SNP hits from GWA.**

**Table S4. Annotation of top SNP hits from aGWA.**

**Table S5. Information on phenotypic and climate traits.**

**Table S6. Correlations between climate and phenotype variables per accession.**

Pearson product-moment correlation coefficients between all phenotype and climate variables of Table S5. Lower triangle shows p-values, upper triangle correlation coefficients. The drought index parameter of choice (m1d_polqua) negatively correlates with the precipitation in the driest month and quarter, bio14 and bio18, respectively.

**Table S7. Correlations between different GWA effects of the 150 polygenic SNPs.**

Pearson product-moment correlation coefficients between SNP effects estimated from GWA of a large subset of all phenotype and climate variables of Table S5.

**Table S8. Canonical Correlation Analysis.**

CCA between GWA effects on different phenotypes and the SNP associations with climate variables.

**Table S9. Polygenic model at different top SNPs groups.**

We applied the Berg & Coop model (66) of polygenic adaptation to different groups of top SNPs and report the value of Q_x_ statistics.

**Table S10. Importance of variables in Random Forest analyses.**

For each random forest model, the importance of bioclimatic variables is reported. For classification random forest, importance is reported as the mean decreased accuracy (MDA) and for regression random forest, importance is reported as the mean square error (MSE). MDA is the number of misclassified observations when removing a variable and MSE is the increase of mean square error produced by removing a variable.

**Table S11. Allele frequency change**

Student’s t-test results of allele frequency changes in the locations of Madrid, Tübingen and Malmö under the three forecasting Genome Environment Models: free migration, principal components control, and geography control.

